# Description of a new genus of primitive ants from Canadian amber, with the study of relationships between stem- and crown-group ants (Hymenoptera: Formicidae)

**DOI:** 10.1101/051367

**Authors:** Leonid H. Borysenko

## Abstract

A detailed study of the holotype of *Sphecomyrma canadensis* Wilson, 1985 from Canadian amber has led to the conclusion that the specimen belongs to a new genus, here named *Boltonimecia* gen.n. Since the taxonomy of stem-group ants is not well understood, in order to find the taxonomic position of this genus, it is necessary to review the classification of stem-group ants in a study of their relation to crown-group ants. In the absence of data for traditional taxonomic approaches, a statistical study was done based on a morphometric analysis of antennae. Scape elongation is believed to play an important role in the evolution of eusociality in ants; however, this hypothesis has never been confirmed statistically. The statistical analysis presented herein lends support to the view that antennal morphology reliably distinguishes stem-group ants from crown-group ants to determine whether a species belongs to one or the other group. This, in turn, may indicate a relationship exists between eusociality and scape elongation. A review of Cretaceous records of ants is made and the higher classification of Formicidae with definitions of stem and crown groups is proposed. Newly obtained data are discussed focusing particularly on the origin, evolution and diversity of ants.

## Introduction

Mammals and birds immediately come to mind when thinking of the groups of animals that began to flourish after the Cretaceous–Paleogene (K-Pg) extinction event. Ants, however, also started to flourish at that time. Ants were rare in the Late Cretaceous, but in the Cenozoic they underwent an explosive radiation to become one of the largest and most widespread groups of terrestrial animals (Hölldobler and Wilson 1990).

Fossilized ants are preserved as imprints in rock or as amber inclusions (reviewed in LaPolla et al. 2013). Among only a handful of amber sites known to contain ants, Canadian amber is of a special interest: dating from a part of the Campanian, 78–79 million years (Ma) old (McKellar and Wolfe 2010), it contains traces of one of the latest Cretaceous ecosystems existed in North America only 10 Ma before the K-Pg extinction event.

It is noteworthy that both crown- (i.e., the descendants of the most recent common ancestor of all extant ants) and stem-group ants (i.e., all taxa that fall outside the crown clade but more closely related to it than to other Aculeata) are found in Canadian amber. These groups had probably coexisted for the most part of the Cretaceous, but only in Canadian amber are they present in almost equal numbers. Four species from Canadian amber represent recent subfamilies - Dolichoderinae (*Eotapinoma macalpini* Dlussky, *Chronomyrmex medicinehatensis* McKellar, Glasier and Engel), Ectatomminae (*Canapone dentata* Dlussky), Aneuretinae (*Cananeuretus occidentalis* Engel and Grimaldi), and at least three species represent the extinct subfamily Sphecomyrminae: *Sphecomyrma canadensis*, *Haidoterminus cippus* McKellar, Glasier and Engel, *Sphecomyrma sp*. (Wilson 1985; Grimaldi et al. 1997; Dlussky 1999a; Engel and Grimaldi 2005; McKellar et al. 2013a, 2013b).

The present paper focuses on the holotype of *Sphecomyrma canadensis.* It has been noted before that the description of *S. canadensis* is unsatisfactory (Dlussky 1996), and the paratype is not conspecific with the holotype (Dlussky 1999a). While examining the holotype, I found that the head, mandibles and antennae’s distal parts were almost invisible. After the amber was polished, some important morphological details were revealed. What had seemed to be a black inclusion hiding the head, turned out to be a thick, raised head platform with posterior stick-like processes. Based on these unique morphological structures, I decided to treat the specimen as belonging to a genus not previously described.

The next step was to find the taxonomic position of this new genus. Needless to say, the classification of stem-group ants is still in its infancy. Their morphological data are scarce, molecular data are impossible to obtain. Stem-group ants have never been the subject of a taxonomic revision, and rarely (Baroni Urbani et al. 1992; Grimaldi et al. 1997; Barden and Grimaldi 2016) were included in a morphological cladistic analysis. At the moment, stem-group ants are assigned to two poorly supported subfamilies, Sphecomyrminae and Armaniinae (Bolton 2003), but some species, such as *Myanmyrma gracilis* Engel and Grimaldi and *Camelomecia janovitzi* Barden and Grimaldi are so bizarre that they cannot be assigned even to these subfamilies and are left as incertae sedis (Engel and Grimaldi 2005; Barden and Grimaldi 2016). There is also a long-standing debate regarding the taxonomic status of Armaniinae, which either represent the most basic stem-group ants (Dlussky 1983), or are sexuals of Sphecomyrminae preserved only in rock imprints due to their large size (Wilson 1987). In the absence of sufficient morphological data, there is a need to invent new methods of taxonomic analysis based on principal differences between stem- and crown-group ants.

In an attempt to fill this gap, I followed the idea that antennal morphology can be a diagnostic tool to distinguish between stem- and crown-group ants. This idea was first expressed by Wilson, Carpenter and Brown (1967) in their diagnosis of Sphecomyrminae, and later explained in terms of evolutionary history and expanded by means of comparative analysis by Dlussky (1983). Since then, antennal characteristics have been used in diagnoses of stem-group subfamilies including Bolton’s system (2003), the most comprehensive for the time being. The fact that they have never been tested by means of statistics is surprising, considering a highly interesting biological background of Dlussky’s hypothesis: scape elongation was necessary for the emergence of eusociality in ants (Dlussky 1983).

The final logical step of the present study was to develop the higher classification of the ants including both stem and crown branches. This important issue that affects our thinking about ant origins has been underestimated in previous works (Ward 2007).

## Materials and Methods

### Examination of the amber inclusion

Photographs were taken with a Nikon D1X digital camera attached to the microscope Leica Z6 APO. The photographs were used to make drawings, which were then computer generated and adjusted using Adobe Photoshop. All measurements were made with an ocular micrometer and are in millimeters. The following measurements were recorded: HL - head length (measured in full-face view as a straight line from the anterior-most point of median clypeal margin to the mid-point of the posterior margin of the head), HW - head width (maximum head width in full-face view), SL - scape length (maximum length excluding articular condyle) F1L-F9L - length of flagellomeres (1^st^ - 9^th^), AL - antenna length, ML - mandible length (maximum length of its horizontal part), WL - Weber’s length (the distance from the anterodorsal margin of the pronotum to the posteroventral margin of the propodeum), TL - total body length.

#### Taxon sampling and morphometry

I used all morphometric data on antennae and heads of stem-group ants available from the literature; also I either made measurements or used published data on antennae and heads of crown-group ants representing all extant subfamilies (Т able S1). For the measurements, only species with both a detailed description in the literature and high resolution images available from AntWeb (Fisher 2002) were selected. I also aimed that the species would cover the broad phylogenetic diversity. All subfamilies in the data set are represented by one species, except for the largest ones (Ponerinae, Dolichoderinae, Formicinae, Myrmicinae), which are represented by two species. Recently, six subfamilies of the dorylomorph group have been subsumed into a single subfamily, Dorylinae (Brady et al. 2014), but here in order to cover broader diversity, all six subfamilies were sampled as valid groups.

Measurements were made on mounted specimens using a Wild M10 stereomicroscope with an accuracy ± 1 µm. For all studied stem- and crown-group ants, I calculated nine indices showing length of antennal parts compared to head length (indices SL/HL, FL/HL, PL/HL, F1L/HL, F2L/HL) and compared to the rest of the antenna (indices SL/FL, PL/(AL-PL), F1L/(AL-F1L), F2L/(AL-F2L)) (Т ables S2, S3). In these indices, HL – head length (see definition above), SL – scape length (see definition above), FL – length of flagellum, PL – length of pedicel, F1L and F2L – length of 1st and 2nd flagellomeres respectively. HL rather than HW was used for the indices, as HL is available for more fossil species; in addition, using HL, the data can be compared with Dlussky’s morphometric data (Dlussky 1983; Dlussky and Fedoseeva 1988). Two more indices, AL/HL and SL/AL (where AL – antenna length, SL – scape length), were only used to compare the data obtained with Dlussky’s data on the Vespoidea and Apoidea (Dlussky and Fedoseeva 1988).

For general observations (the shape of the pedicel, the morphology of middle and terminal antennal segments), high resolution images available from AntWeb were used. To study the width of the petiolar attachment of the Armaniinae, the petiolar index, PG/PH (where PG - the width of the petiole in the broadest point of its attachment to the gaster; PH - the maximum height of the petiole) was calculated.

#### Statistical analysis

Two statistical tests for equality of means were performed: the Student’s t-test and a one-way ANOVA with planned comparisons. In addition, correlation and regression analysis as well as canonical discriminant analysis were performed.

Power analysis was carried out using the program G*Power (Faul et al. 2007). Despite the scarcity of the data on extinct taxa, in all cases where the t test showed statistically significant results, statistical power was also high. An unbalanced design in an ANOVA, however, could be a problem (McDonald 2014). Indeed, the results showed that the unbalanced design in which extinct ants are underrepresented, had low statistical power. In such a case, the only way to confirm reliability of the results is to reduce large groups to a size of the smallest one, and run the power analysis and ANOVA again. After such manipulation, high statistical power was obtained, whereas the ANOVA results were almost identical to those of the unbalanced design.

The next concern about the reliability of the statistical results was a measurement error. Since the measurements of extinct taxa were taken independently by different persons, with different material and calibration, those slight differences might presumably alter the conclusions obtained. In order to check the extent to which the results were sensitive to errors, 10% (a large measurement error) were added/subtracted to/from the indices and to/from the measurements. A random number generator was used to apply these modifications. Then the modified data set was analyzed with the ANOVA and t test again. In all cases no noticeable effect was observed, so the statistical model proved to be robust and not sensitive to large errors.

Supplementary material (Tables S1-S18) available at https://goo.gl/3MWImf

The collection acronym used in this study is as follows: CNC - Canadian National Collection of Insects, Arachnids and Nematodes (Ottawa, Canada).

### Systematic Palaeontology

**Family Formicidae Latreille, 1809**

**Subfamily Sphecomyrminae Wilson and Brown, 1967**

**Tribe Boltonimeciini, trib.n.**

**Genera.** *Boltonimecia* gen.n. (type genus), *Zigrasimecia* Barden and Grimaldi, 2013.

**Diagnosis (workers)**. See “The higher classification of the ants” below.

Genus *Boltonimecia*, gen.n.

**Type and only known species.** *Boltonimecia canadensis* (Wilson, 1985)

**Diagnosis**. As for the tribe.

**Etymology**. This genus is dedicated to the renowned myrmecologist Barry Bolton.

*Boltonimecia canadensis* **(Wilson, 1985), comb.n.**

Fig. 1, 2

**Figure 1.**
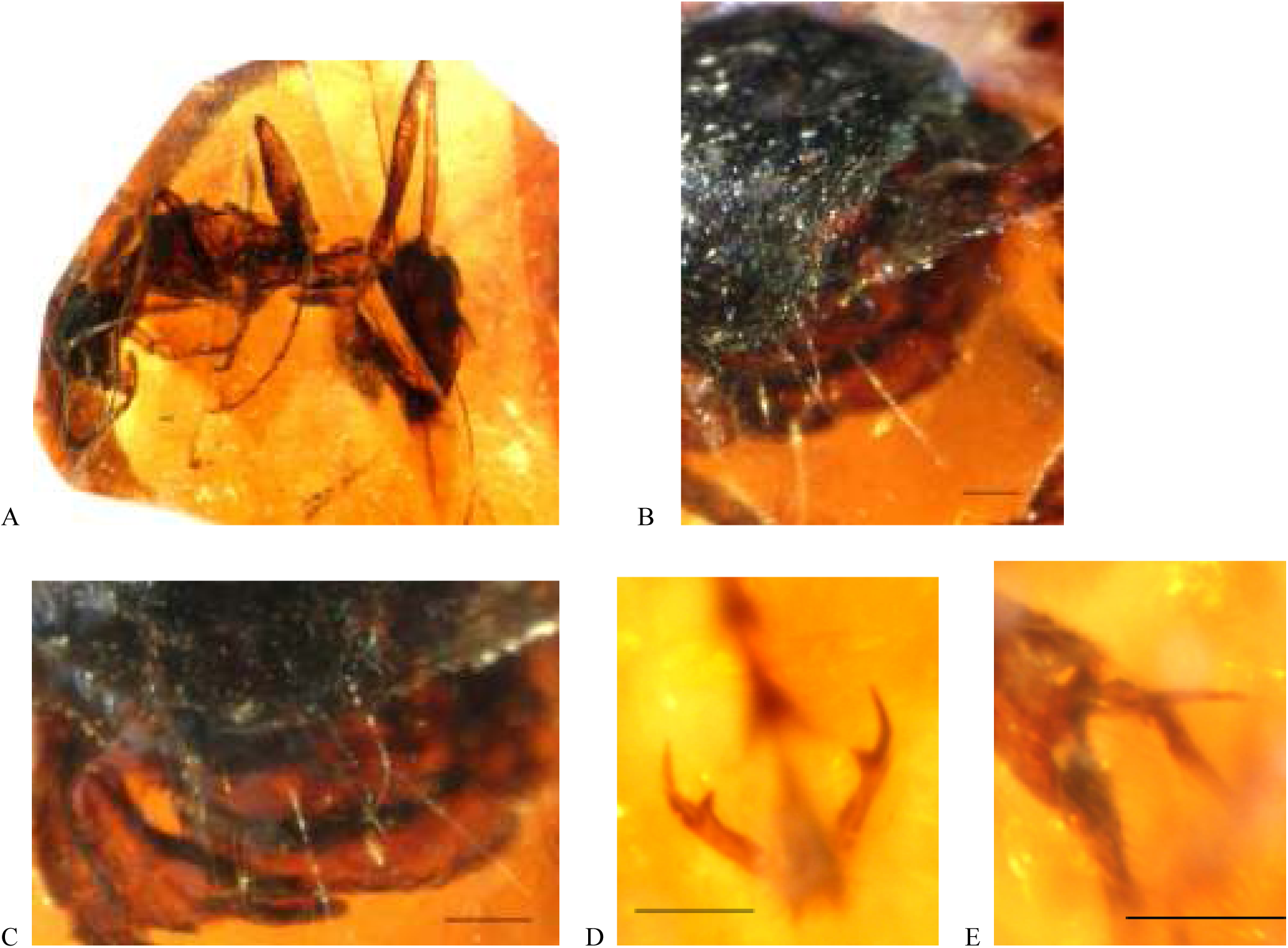
Photographs of *Boltonimecia canadensis.* **A)** General habitus, lateral view. **B)** Part of head, anterodorsal view. **C)** Clypeus and mandibles. **D)** Pretarsal claws. **E)** Metatibial spurs. Scale line = 0.1 mm.

**Figure 2.**
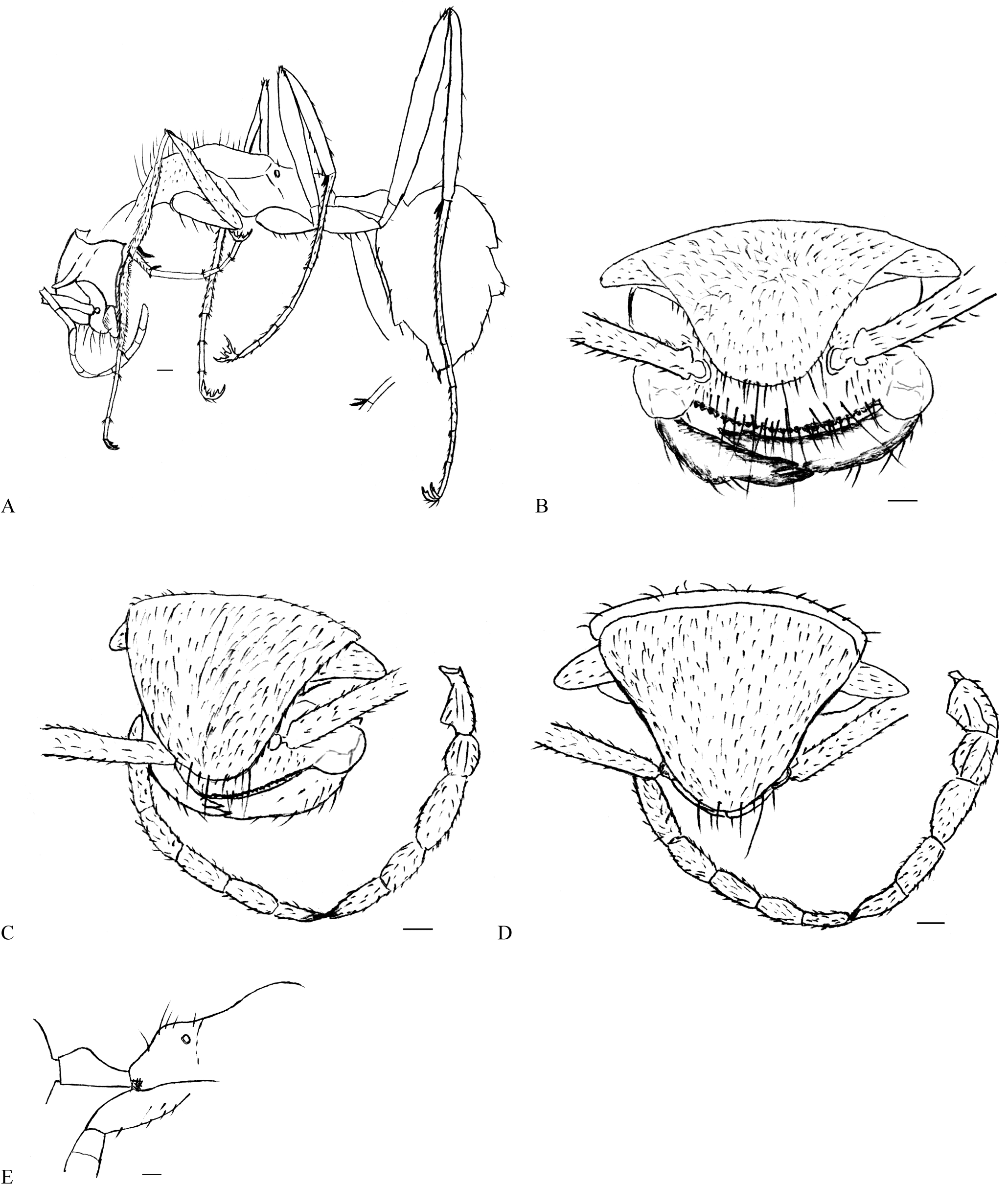
Drawings of *Boltonimecia canadensis*. **A)** General habitus, lateral view (pubescence on mesosoma, legs, and gaster omitted). **B)** Head, anterodorsal view (reconstruction). **C)** Head, anterolateral view. **D)** Head, dorsal view (reconstruction). **E)** Propodeum, lateral view. Scale line = 0.1 mm.

#### Taxonomic history

*Sphecomyrma canadensis* Wilson, 1985: 206, fig. 1, 2 (w.)

#### Materials examined

Holotype, worker. The specimen is preserved in a clear orange piece of Canadian amber, 8×3×2 mm in size, from Medicine Hat, Alberta (J. F. McAlpine, CAS 330), held in the CNC. The preservation is excellent, except for the mesosoma, left flagellum and the right side of the head, which are flattened due to compression; also a small proximal part of both scapes is gone.

The specimen described as the paratype of *Sphecomyrma canadensis* (Wilson, 1985) (CAS 205, CNC) is represented by poorly preserved body fragments without taxonomically informative characters, and thus should not be regarded as the paratype.

#### Diagnosis

As for the genus.

#### Redescription of type

HL 0.73 mm; HW 0.8 mm; SL 0.5 mm; ML 0.48 mm; WL 1.2 mm; TL 3.4 mm.

Head small compared with the rest of the body (1/5 of body length), slightly wider than long, seems to be triangular when seen from above. Its dorsal part thick, raised and curved in profile, thus forming a “shield” (Fig. 2B-D). Two stick-like processes directed anterolaterally protrude from the posterolateral edges of the head. Compound eyes and ocelli absent. The stick-like processes are doubtfully deformed eyes, since they have no trace of visible facets and are densely covered with appressed pubescence, similar to that on the front of the head. Clypeus small, in profile strongly convex, does not extend back between the antennal sockets; its lateral margins produced as semicircular lobes covering the insertions of the mandibles; its anterior margin bears 25 peg-like setae 0.01 mm long. Clypeal width 0.5 mm (without lateral lobes), length - 0.15 mm.

Mandibles linear, two toothed, curved at almost 90 °. The apical tooth is longer than the preapical one: 0.15 mm and 0.06 mm, respectively. On the inner side of the apical tooth there is a longitudinal impression, which probably corresponds to the position of the other mandible when mandibles are closed. Mandible length: 0.48 mm (horizontal part), 0.17 mm (vertical part).

Antennae 11-segmented; when laid back the apex of the scape just reaches the occipital margin. Insertions of antennae are not far apart (0.17 mm), partly exposed, and touching the clypeus. Toruli not fused to the frontal lobes. Antennal segment measurements (mm): SL - 0.5, PL - 0.2, F1L - 0.15, F2L - 0.2, F3L-F8L - 0.17, F9L - 0.25; AL - 2.32.

Metanotal groove distinct. Propodeum slightly lower than promesonotum, without teeth or spines; propodeal spiracles high. Orifice of metapleural gland protected by guard setae. Petiole 0.4 mm long, pedunculate (peduncle 0.1 mm long, node 0.2 mm high). Gaster ovate, 1.1 mm long; helcium projects from abdominal segment III low down on its anterior face; abdominal segment IV without presclerites. Sting present; length of its extruded portion 0.05 mm.

Legs long: 3.3 mm forelegs (shortest), 5.07 mm hind legs (longest; 1.5 times of body length); measurements of leg segments are given in Table 1. Pretarsal claws with one preapical tooth (Fig. 1D). Each tarsal segment with two stiff setae on both sides apically; protibia with one pectinate and two simple spurs; mesotibia and metatibia with one pectinate and one simple spur (Fig. 1E).

**Table 1.** Leg segment measurements (in mm) of *Boltonimecia canadensis*.

| Segment | Front leg | Middle leg | Hind leg |
| --- | --- | --- | --- |
| coxa | 0.40 | 0.34 | 0.50 |
| femur | 0.75 | 0.85 | 1.25 |
| tibia | 0.75 | 0.75 | 1.20 |
| tarsal segment 1 | 0.50 | 0.60 | 0.90 |
| tarsal segment 2 | 0.15 | 0.25 | 0.25 |
| tarsal segment 3 | 0.20 | 0.20 | 0.25 |
| tarsal segment 4 | 0.10 | 0.15 | 0.17 |
| tarsal segment 5 | 0.15 | 0.15 | 0.25 |
| pretarsal claws | 0.15 | 0.15 | 0.15 |
| *total length* | *3.30* | *3.60* | *5.07* |

Head dorsum, antennae, stick-like processes covered with dense short appressed pubescence. Lateral head margins with erect and suberect hairs. Anterior clypeal margin, middle of clypeus, anterior margin of the “shield” bear long erect hairs directed anteriorly. External mandibular margins covered with suberect hairs. Mesosoma, legs, and gaster completely covered with short appressed pubescence. In addition to pubescence, middle and hind tibiae covered with sparse suberect hairs; erect hairs project from the apex of middle and hind femurs; ventral surfaces of coxae with long erect hairs; pronotum and propodeum bear long white erect hairs tapering to sharp points (especially long on pronotum); gaster covered with sparse suberect hairs, which is longer on the ventral surface.

Body sculpture not distinct. Body colored as surrounding amber material, but the ventral surfaces of coxae, gaster, proximal halves of tibiae, lower half of propodeum are brown to black. Head dorsum and pronotum black, opaque.

Gyne and male unknown.

#### Discussion

Some authors have already noticed that *Sphecomyrma canadensis* shares no synapomorphy with *Sphecomyrma* Wilson and Brown, namely, it lacks an elongated F1 (Dlussky 1996; Grimaldi et al. 1997; Engel and Grimaldi 2005). Dlussky and Fedoseeva have even suggested that because of unique antennal characters, such as the elongated scape and pedicel, this species should be assigned to its own new subfamily (Dlussky and Fedoseeva 1988). Here more unique characters that set *B. canadensis* apart from all known Cretaceous ants were discovered.

*Boltonimecia canadensis* has 11-segmented antennae - a unique character unknown in any other Cretaceous ants. A small proximal part of both scapes is missing, but the left scape-pedicel joint is intact, and thus one can confidently infer about the scape length. This joint is not the one between the pedicel and F1 because (1) it is curved near the base, which is a distinct feature of the pedicel only; and (2) spatial localization of the scape and the flagellum infers that there is no room left for one more antennomere between them. In general, the relative size of the antennomeres is not different from Wilson’s drawings (Wilson 1985). The only exception is the distal parts of the antennae, which are curled back under the head and were almost invisible before additional polishing; this probably led to Wilson’s assumption about the 12-segmented antenna as in *Sphecomyrma*.

The most distinctive character of *B. canadensis* is the specialized head: the thick “shield” (most likely formed by the laterally expanded frontal lobes), stick-like processes, and long sensory hairs anteriorly. Some Myrmicinae, *Aulacopone relicta* Arnol’di (Heteroponerinae) and the Agroecomyrmecinae also have shield-like heads formed by the expanded frontal lobes. In the Myrmicinae, unlike *B. canadensis,* the median portion of the clypeus usually inserted between the antennal sockets (Bolton 2003). In *A. relicta,* the clypeus is shallow, but, unlike *B. canadensis,* the antennal insertions are close together, and the fronto-clypeal part of the head is extended forwards and covers the mandibles (Taylor 1980). All species of an enigmatic subfamily Agroecomyrmecinae have shield-like heads; arboreal *Ankylomyrma coronacantha* Bolton also has dentiform processes on the occipital margin, projecting especially strongly at the occipital corners (Bolton 1973), in which it closely resembles the specialized head of *Boltonimecia*. However, in the Agroecomyrmecinae the clypeus is large and broadly inserted between the frontal lobes. In most Ponerini the clypeus is narrowly inserted between the antennal sockets, but their confluent frontal lobes are thin and never form a “shield”. In the Proceratiinae, the clypeus is reduced, and the antennae inserted close to the anterior head margin. Some Proceratiinae (*Discothyrea* Roger) have the frontal lobes fused and forming a small raised platform behind the level of the antennal sockets, the sides of which are strongly convergent anteriorly (Bolton 2003).

Unfortunately, all these interesting morphological similarities tell us little about the evolutionary relationship of *B. canadensis*, because similarities between a stem-group species and a living species have likely evolved through parallel evolution as a result of adaptation to similar habitats. Regarding the last, the lack of eyes and ocelli, as well as long sensory hairs on the anterior head margin may suggest a cryptic lifestyle. However, blind extant ants are not always exclusively subterranean (e.g., *Dorylus* Fabricius), and long legs of *Boltonimecia* speak in favor of an arboreal or above-ground lifestyle. The possibility that the eyes of *Boltonimecia* are reduced to a single facet, and thus simply invisible in amber cannot be rejected either. Therefore, determining the evolutionary relationship and the phylogenetic position of *B. canadensis* is a major challenge. In order to address this issue, it is necessary first to consider the classification of stem-group ants and their relationship with crown-group ants - a task that has rarely been undertaken to date.

### What is the difference between stem- and crown-group ants?

The stem- and crown-group concepts have been developed with the introduction of phylogenetic systematics in the mid-20th century (Hennig 1966). By definition, the crown-group is the clade that includes the last common ancestor of all living taxa and all its descendants; the pan- or total group is the clade comprised of the crown-group and all taxa more closely related to it than to any other extant organism; finally, the stem-group is a paraphyletic group composed of the total group excluding the crown-group (e.g., Ax 1985). From this, the ant subfamilies Sphecomyrminae and Armaniinae, as well as incertae sedis genera *Myanmyrma* Engel and Grimaldi and *Camelomecia* Barden and Grimaldi are stem-group ants; all living ant taxa plus most of the known extinct genera, such as *Kyromyrma* Grimaldi and Agosti and *Canapone* Dlussky, are crown-group ants; both groups, together, form the pan-group Formicidae, the ants.

Among the plesiomorphic characters useful to differentiate between stem- and crown-group ants are the following: the broad attachment of the petiole to the gaster, pretarsal claws with a preapical tooth, bidentate mandibles, two spurs on meso- and metatibiae, short scape, trochantellus, peg-like setae on the anterior margin of the clypeus, and ocelli (Dlussky 1983, Bolton 2003, LaPolla et al. 2013). At first glance some of these characters seem to be important but others are not that reliable.

The petiole in the non-ant Vespoidea is less differentiated and more broadly attached to the gaster than in the Formicidae, except for the only ant subfamily Amblyoponinae. This may be a good indication that the broad attachment is a clearly plesiomorphic condition. If this assumption is correct, all presently known stem-group ants should be considered as quite advanced, because they all have differentiated petioles, similarly to crown-group ants. The problem of the broad petiole-gaster attachment in the Armaniinae (Dlussky 1983) remains open, and will be discussed in details below.

A preapical tooth on the pretarsal claws is present in many crown-group ants, such as the poneroids and primitive formicoids - Myrmeciinae, Pseudomyrmecinae, Dorylinae (Dlussky and Fedoseeva 1988), as well as in virtually all stem-group ants. It is also common in other Vespoidea.

Bidentate mandibles are common in the Vespoidea and Apoidea, and universal for stem-group ants. In crown-group ant females, bidentate mandibles are mainly present in the poneroids and primitive formicoids, while being quite rare in advanced formicoids - Formicinae, Myrmicinae, Dolichoderinae (Bolton 2003). Similarly, in crown-group ant males, they are common in the poneroids, but rare in the formicoids: in the Formicinae and Myrmicinae they are present in 1/4 of the genera; rarely present in the Dolichoderinae, ectaheteromorphs, Pseudomyrmecinae (Bolton 2003). It is believed that bidentate mandibles evolved in crown-group ants as a result of the reduction of triangular mandibles (Dlussky and Fedoseeva 1988).

Two spurs on meso- and metatibiae are common in the poneroids and primitive formicoids (Bolton 2003), in all stem-group ants, as well as in other Vespoidea. *Haidoterminus cippus* was described with a single metatibial spur and two mesotibial spurs (McKellar et al. 2013b), which most probably is the result of poor preservation of the legs, because this feature is not known in other Formicidae.

The trochantellus is absent in crown-group ants, except for a putative crown-group species *Cananeuretus occidentalis* (Engel and Grimaldi 2005). In stem-group ants, it is present in one species of *Haidomyrmex* (*H. scimitarus* Barden and Grimaldi) (Barden and Grimaldi 2012), *Haidoterminus cippus* (McKellar et al. 2013b), both species of *Zigrasimecia* Barden and Grimaldi (Barden and Grimaldi 2013; Perrichot 2014), some species of *Gerontoformica* Nel and Perrault (Barden and Grimaldi 2014), and also in the males of *Baikuris* Dlussky (Dlussky 1987; Grimaldi et al. 1997; Perrichot 2015). In the Armaniinae, this character is not obvious: *Armania* Dlussky and *Pseudarmania* Dlussky have been reported either with or without trochantelli (Dlussky 1983; Wilson 1987; Dlussky and Fedoseeva 1988).

Peg-like setae on the anterior clypeal margin are thought to be an important plesiomorphy for the Formicidae (Engel and Grimaldi 2005). In stem-group ants, this character is present in *Boltonimecia*, *Gerontoformica*, *Myanmyrma*, *Zigrasimecia*; in crown-group ants, it is found in the subfamily Amblyoponinae. Outside ants, peg-like setae exist in some Tiphiidae (*Myzinum* Latreille) and Bradynobaenidae (*Apterogyna* Latreille).

Ocelli in workers are often considered an ant plesiomorphy (Engel and Grimaldi 2005). In crown-group ants, this character is common in extant taxa, mainly in the formicoid clade (subfamilies Myrmeciinae, Pseudomyrmecinae, Dolichoderinae, Formicinae, some Dorylinae), but absent in such Cretaceous genera as *Eotapinoma*, *Chronomyrmex*, and *Kyromyrma*. In stem-group ants, ocelli are present in some Sphecomyrmini.

To sum up, all the mentioned characters can hardly be used to differentiate reliably between stem- and crown-group ants. As noted by Dlussky (1983), the most reliable character is probably the elongated scape of crown-group ants, which allows brood and food manipulation and thus favors the emergence of eusociality. It is now time to examine this character, along with other morphometric features of the ant antenna, in detail.

### The antennal structure as a key distinction between stem- and crown-group ants

Scape length of more than 25% of antennal length is thought to be a characteristic of extant (i.e., crown-group) ants (Dlussky and Fedoseeva 1988), although the diagnosis of Sphecomyrminae (Bolton 2003) stated that a “short scape” of Sphecomyrminae means “0.25 times length of funiculus”. The role of other antennal parts in distinguishing stem-group ants from crown-group ants may be no less important.

The pedicel of all insects contains the Johnston’s organ - a mechanosensory chordotonal organ responsible for hearing, graviception (Kamikouchi et al. 2009) and electric field changes, which may play a role in social communication (Greggers et al. 2013). According to Dlussky and Fedoseeva (1988), the pedicel in crown-group ants is shorter than in stem-group ants, it is narrowed and curved at the base. This enables close contact of the pedicel and the scape, resulting in greater freedom and accuracy of movement of the flagellum, which together with scape elongation led to the emergence of eusociality in ants (Dlussky and Fedoseeva 1988).

The first flagellomere is the longest flagellar segment in stem-group ants, the males of primitive crown-group ants (Dlussky 1983), and the Aculeata closely related to the ants (Engel and Grimaldi 2005); so it is a symplesiomorphic character (Engel and Grimaldi 2005). Bolton (2003) mentioned this character as a synapomorphy of the Sphecomyrmini (while the longest flagellar segment in the Haidomyrmecini is the second one). In crown-group ants, the first and second flagellomeres are not different from other flagellomeres, except for the elongated terminal one (Dlussky and Fedoseeva 1988).

In stem-group ants, segments beyond the second flagellomere decrease in length towards the apex of the antenna, while in crown-group ants they often increase, ending in a club-shaped expansion of terminal segments (Dlussky and Fedoseeva 1988). In crown-group ant females, the club is common in advanced taxa, except for the formicines in which 3/4 of the genera lack it. Crown-group ant males also rarely have clubs (Bolton 2003). Finally, the entire flagellum in stem-group ants is long and flexuous (Bolton 2003).

Making his hypothesis from a comparison of ants with other Aculeata, Dlussky, however, has not provided any statistical support. This has resulted in criticism and even removal of a character “short scape” from the data matrix as it is “difficult to define” (Grimaldi et al. 1997). Below I check Dlussky’s hypothesis using a statistical analysis of antennal indices and try to expand and generalize the aforementioned observations on the antennal structure.

#### A comparison of antennal indices of crown- and stem-group ants

Although the indices of the Cretaceous ant males (since none of the Cretaceous ant males has been explicitly associated with conspecific stem-group ant females, they are not called “stem-group ant males” throughout the paper) are within the range of the indices of the crown-group ant males, in most cases they are shifted from the crown-group male mean (Table S3). The statistical tests showed that some differences between these indices were significant (Table S15):

(1) Scape. The indices SL/HL did not show statistically significant differences, while for SL/FL such a difference existed. The latter can be explained by a considerably longer flagellum of the Cretaceous ant males.
(2) Flagellum. The mean of FL/HL is noticeably greater in the Cretaceous ant males than in the crown-group ant males, although a P-value is quite high.
(3) Pedicel. For PL/HL, the difference was statistically insignificant, while for PL/(AL-PL), it was on the verge of significance. The latter again results from a longer flagellum of the Cretaceous ant males.
(4) The first and second segments of the flagellum. The means of F1L/HL and F2L/HL are noticeably greater in the Cretaceous ant males, although P-values are quite high. The differences between F1L/(AL-F1L) as well as F2L/(AL-F2L) are not well understood due to low statistical power.

The male regression lines were similar for FL/HL, F1L/HL, and F2L/HL (Fig. 3).

**Figure 3.**
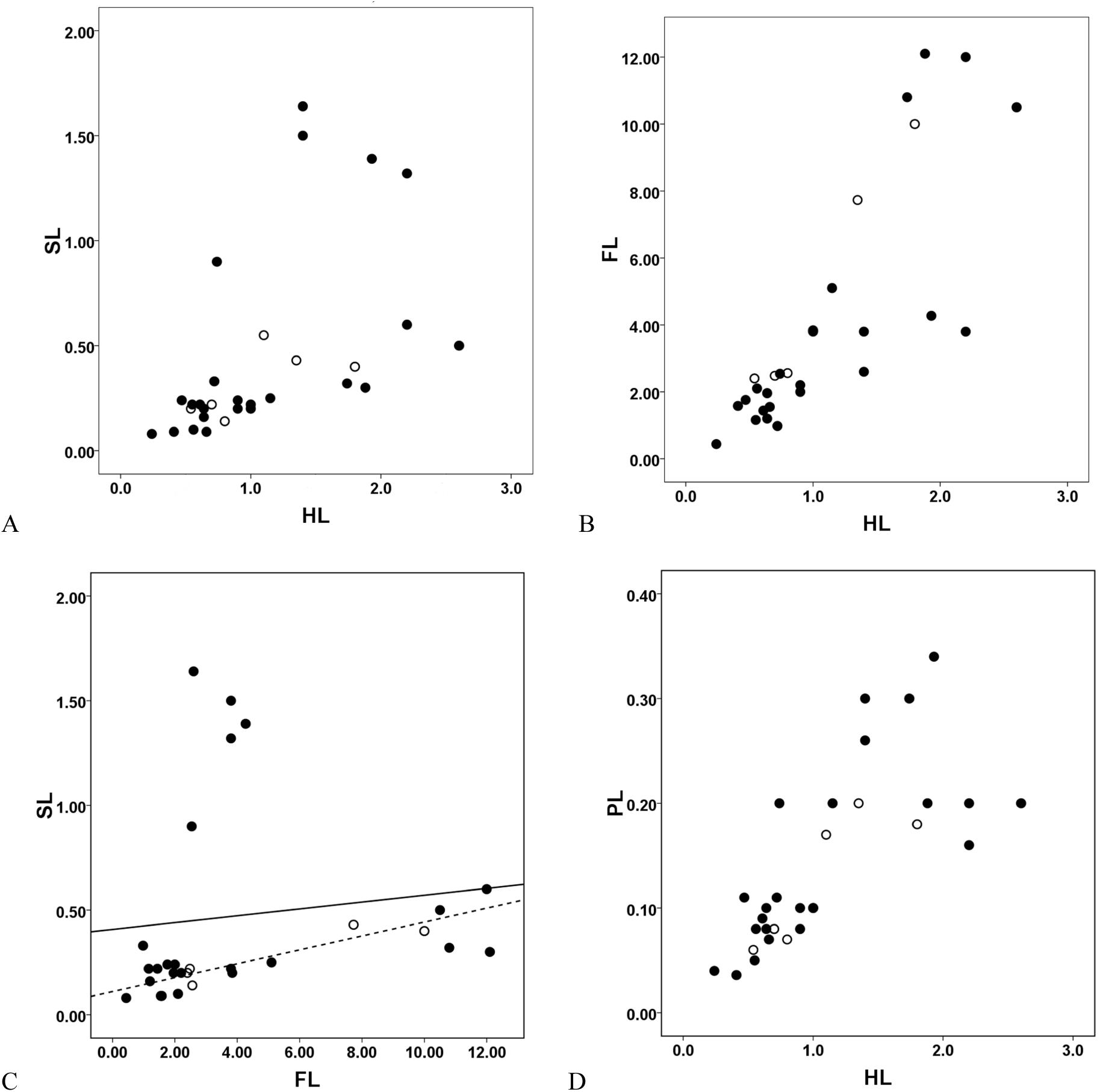
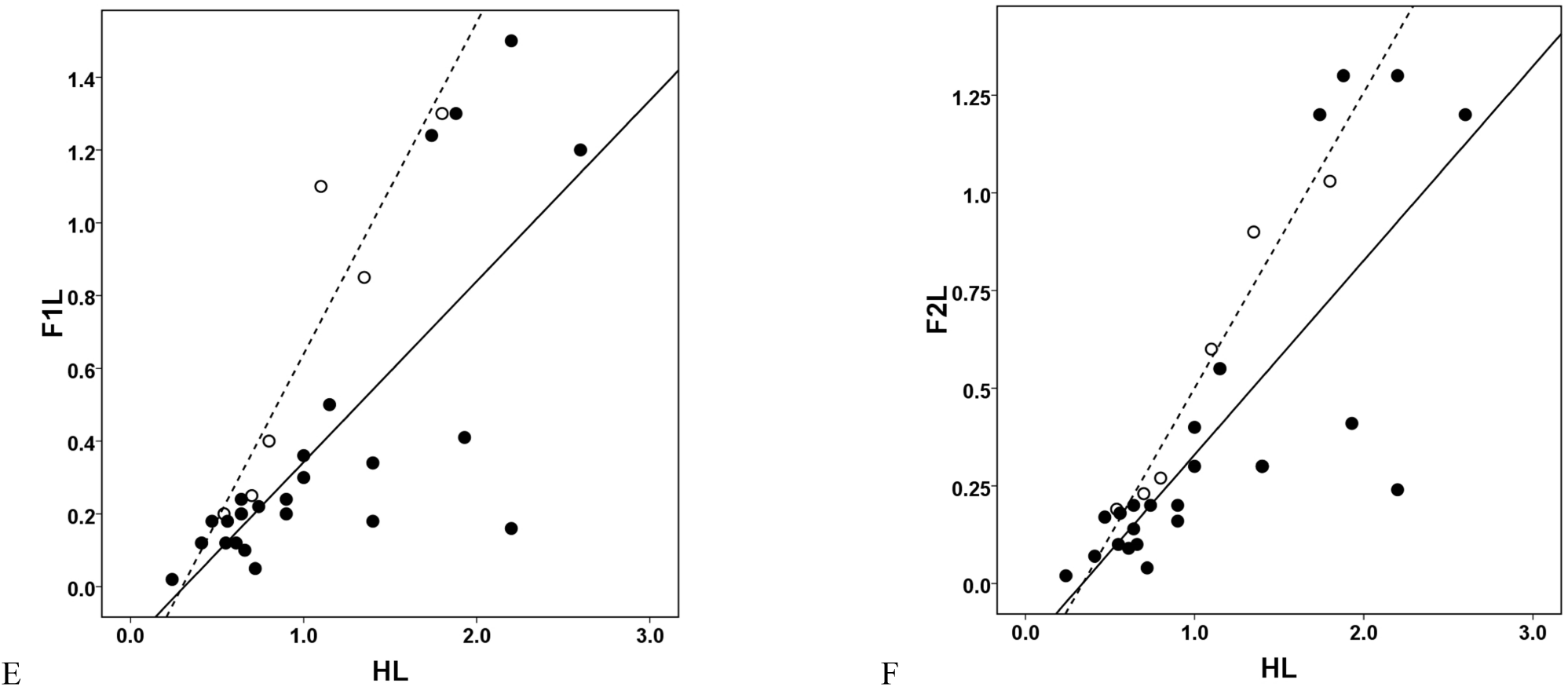
Bivariate plots (males). **A)** Scape length versus head length. **B)** Flagellum length versus head length. **C)** Scape length versus flagellum length. **D)** Pedicel length versus head length. **E)** Flagellomere 1 length versus head length. **F)** Flagellomere 2 length versus head length. (Note: Filled circles are crown-group ants; open circles are stem-group ants; regression lines for both groups (crown-group ants - solid line, stem-group ants - dashed line) are shown in cases where statistical difference between them was found).

The low sample size of the Cretaceous ant males prevents any firm conclusions being drawn; the available data, however, suggest that the Cretaceous ant males might have a longer flagellum, F1, and F2, compared to the crown-group ant males.

The situation is completely different for females, as the statistical analysis showed highly significant differences between the crown- and stem-group ants. The ANOVA analysis for five groups (extant crown-group ants, Cretaceous crown-group ants, Sphecomyrmini, Haidomyrmecini, Armaniinae) showed that the means of all the indices differed significantly, except for the pedicellar indices: SL/HL: F_4,79_ = 18.38, P < 0.0001; FL/HL: F_3,74_ = 23.29, P < 0.0001; PL/HL: F_4,78_ = 1.47, P = 0.22; F1L/HL: F_4,78_ = 34.36, P < 0.0001; F2L/HL: F_4,78_ = 18.01, P < 0.0001; SL/FL: F_3,74_ = 87.78, P < 0.0001; PL/(AL-PL): F_3,74_ = 0.35, P = 0.79; F1L/(AL-F1L): F_3,74_ = 38.58, P < 0.0001; F2L/(AL-F2L): F_3,74_ = 18.07, P < 0.0001. A planned comparison revealed the following picture.

All indices of the Cretaceous crown-group ants are close to the means of the extant crown-group ants (Table S2); the statistical analysis showed no differences between the two groups (Tables S4-S12). The relationships among other groups were more complicated.

Scape (indices SL/FL, SL/HL):

(1) For SL/FL, the stem-group ants differed significantly from the crown-group ants in having lower mean values. The Haidomyrmecini were significantly different from both the crown-group ants and Sphecomyrmini (Table S9).
(2) For SL/HL, the Sphecomyrmini and Armaniinae have significantly lower mean values than the crown-group ants. The Haidomyrmecini have greater means, which are intermediate between the means of the crown-group ants and Sphecomyrmini, Armaniinae (Table S4); Haidomyrmecini’s values are often seen in the crown-group ants (Table S2).
(3) For SL/HL, the Armaniinae were not different from the Sphecomyrminae and Sphecomyrmini (Table S4); for SL/FL, the only available index of the Armaniinae is close to the mean of the Sphecomyrmini (Tables S2, S4).
(4) *Myanmyrma*’s indices lie close to the regression line of the Sphecomyrminae (Fig. 4A, 4C). *Myanmyrma*’s SL/HL is similar to the mean of the Sphecomyrmini, Armaniinae, and the lowest value of the crown-group ants obtained in this study, the index of *Pseudomyrmex pallidus* Smith. *Myanmyrma*’s SL/FL is the lowest one, but it is quite close to the minimum value of the Sphecomyrmini (index of *Gerontoformica contegus* Barden and Grimaldi) (Table S2).
(5) *Boltonimecia*’s SL/HL is close to the mean of the crown-group ants (similar indices are seen in the Dorylinae, Proceratiinae, Myrmicinae, Ponerinae, Agroecomyrmecinae), but greater than that of all Sphecomyrmini and Haidomyrmecini (except for *Haidoterminus cippus*). *Boltonimecia*’s SL/FL is greater than that of most Sphecomyrmini, but lower than that of several species of the Haidomyrmecini (Table S2).

**Figure 4.**
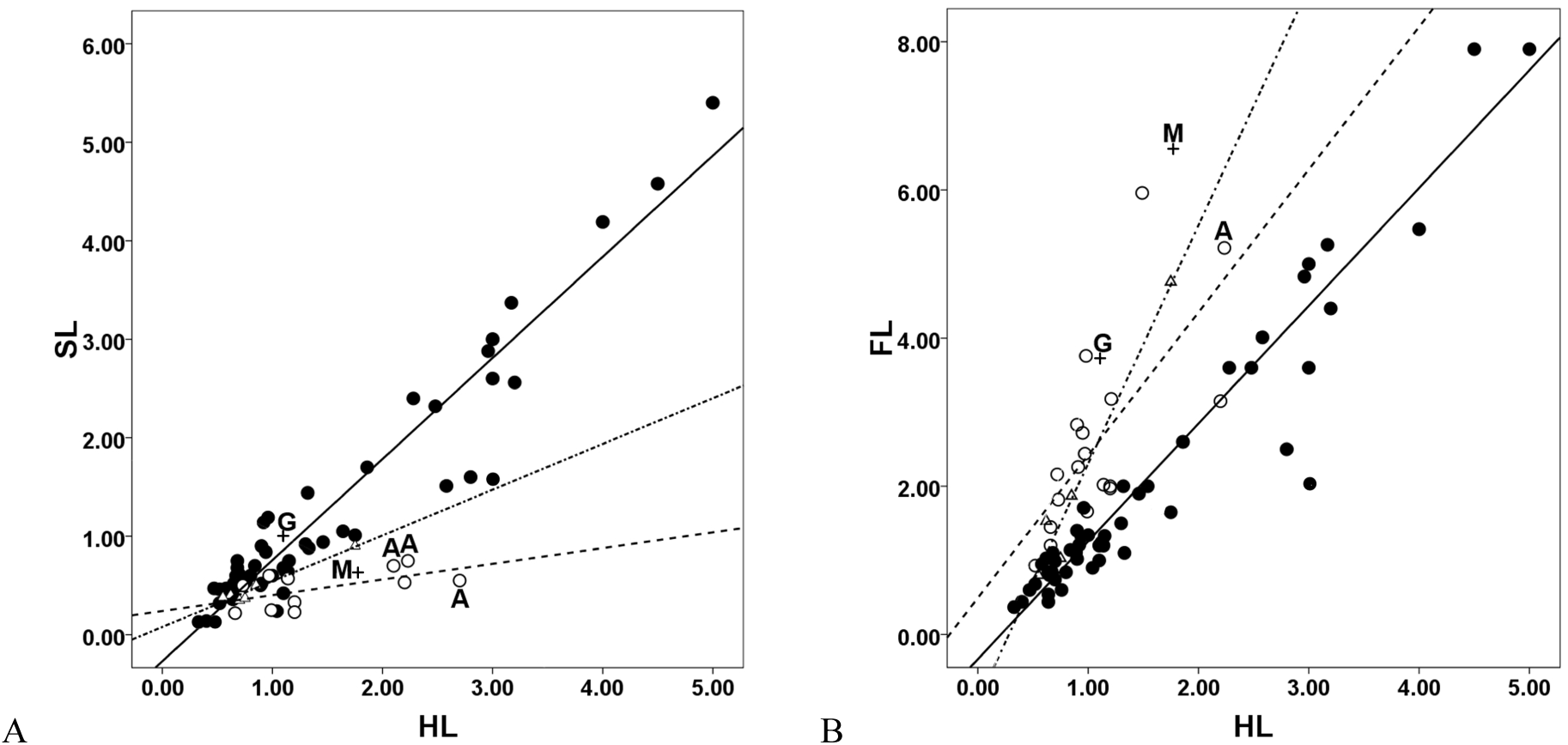
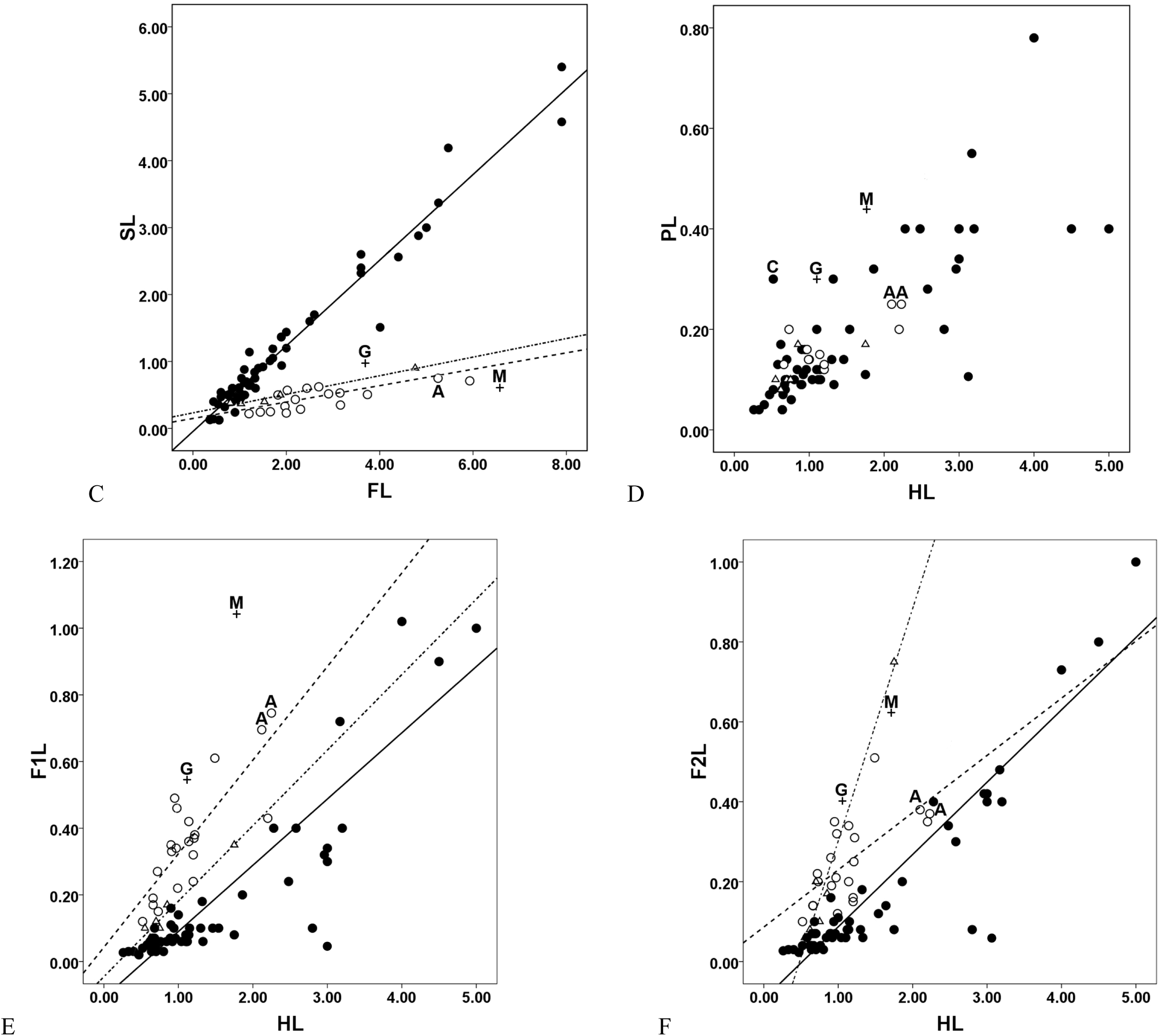
Bivariate plots (females). **A)** Scape length versus head length. **B)** Flagellum length versus head length. **C)** Scape length versus flagellum length. **D)** Pedicel length versus head length. **E)** Flagellomere 1 length versus head length. **F)** Flagellomere 2 length versus head length. (Note: Filled circles are crown-group ants; open circles are Sphecomyrmini; triangles are Haidomyrmecini; A - Armaniinae (sensu Bolton 2003); C - Cananeuretus occidentalis; G - *Gerontoformica cretacica*; M - *Myanmyrma gracilis*. Regression lines for crown-group ants (solid line), Sphecomyrmini (dashed line), and Haidomyrmecini (dash-dotted line) are shown in cases where statistical difference between at least two groups was found).

Pedicel (indices PL/(AL-PL), PL/HL):

1. For both indices, there was no statistical difference between the groups studied (Tables S6, S10; Fig. 4D). Such stability, as already noted, may be explained by an important function of the pedicel.
2. The greatest PL/HL is seen in *Cananeuretus occidentalis*, followed by *Gerontoformica cretacica* Nel and Perrault, *Boltonimecia canadensis*, *G. rugosus* Barden and Grimaldi and *Myanmyrma gracilis*; all four indices are close to one another and to the greatest value among the crown-group ants found in *Martialis heureka* Rabeling and Verhaagh (Table S2). *G. cretacica*, *M. gracilis* and *C. occidentalis* appear separate from the rest of the species in the bivariate plot (Fig. 4D).

Flagellum (index FL/HL):

(1) The stem-group ants were always statistically different from the crown-group ants in having longer flagella (Table S5).
(2) The greatest FL/HL (i.e., the longest flagellum) is in *Gerontoformica rubustus* Barden and Grimaldi*, G. magnus* Barden and Grimaldi*, Myanmyrma gracilis* and *G. cretacica* (Table S2). The extremely elongated flagella of these species resemble only the male flagella (Table S3).
The Haidomyrmecini were not different from the Sphecomyrminae (Table S5).
The indices of *Boltonimecia canadensis* and the only species of the Armaniinae for which FL/HL can be calculated (*Pseudarmania rasnitsyni* Dlussky) are about equal to the mean of the Sphecomyrminae (Table S2).

The first segment of the flagellum (indices F1L/(AL-F1L), F1L/HL):

(1) Most stem-group ants, except for the Haidomyrmecini, had significantly greater means than the crown-group ants (Tables S7, S11). The Haidomyrmecini’s means are intermediate between those of the Sphecomyrmini and crown-group ants, with weak statistical relationships (Tables S7, S11).
(2) The Armaniinae were not statistically different from the Sphecomyrminae and Sphecomyrmini (Table S7).
(3) The indices of *Boltonimecia* are within the range of the indices of the Sphecomyrminae, although the former are not as great as the indices of most Sphecomyrmini (Table S2).
(4) *Myanmyrma*’s F1L/HL is the greatest of all the species reported, followed by *Gerontoformica subcuspis* Barden and Grimaldi, and *G. cretacica* (Table S2); *Myanmyrma* is separate from the rest of the species in the bivariate plot (Fig. 4E). *M. gracilis* and *G*. *subcuspis* have the greatest F1L/(AL-F1L) (Table S2).

The second segment of the flagellum (indices F2L/(AL-F2L), F2L/HL):

(1) The indices of the stem-group ants are usually significantly greater than those of the crown-group ants (Tables S8, S12).
(2) In the bivariate plot of F2L vs HL (like in the plots of SL vs HL and PL vs HL) the Armaniinae lie close to *Sphecomyrma mesaki* Engel and Grimaldi (Fig. 4A, 4D, 4F).
(3) *Boltonimecia*’s indices are close to the mean of the Sphecomyrminae (Tables S2, S8, S12).
(4) F2L/HL of *Myanmyrma gracilis*, *Gerontoformica subcuspis*, and *G. cretacica* is close to the greatest value found in *Haidomyrmex scimitarus*. The greatest F2L/(AL-F2L) is found in *H. scimitarus* and *G*. *subcuspis* (Table S2).

#### General observations on middle and terminal flagellomeres

(1) Crown-group ant females: following F2 or F3, segment width (and often length) increases. This increase may be gradual or sharp (forming an antennal club). A slight width increase is seen even in species with filiform antennae (marked “0” in Appendix 2 of Bolton 2003), for example in the Paraponerinae, Formicinae, Myrmeciinae. Also in the species with filiform antennae, the terminal flagellomere is often about 1.5 times longer than the penultimate one. The exceptions are the genera without an increase in width (*Eciton* Latreille) or length (*Paraponera* Smith). In some cases, the pattern resembles that of some stem-group ant females: segment length diminishes towards the apex (*Eciton*, *Nomamyrmex* Borgmeier, *Leptomyrmex* Mayr), but often such a decrease is accompanied by the elongation of the terminal segment (*Eciton, Nomamyrmex, Leptomyrmex*), and/or with a slight increase in width of last segments (*Myrmecia* Fabricius).

(2) Stem-group ant females: Antennae are always filiform without a club; middle and terminal segments are similar in size, but in some cases the most distal segments are slightly thicker (in *Gerontoformica orientalis* Engel and Grimaldi, *G. contegus*, *Haidomyrmodes mammuthus, Haidoterminus cippus*). The terminal segment is often about 1.2-1.5 times longer than the penultimate one (except *G. cretacica*, *Haidomyrmex scimitarus*, *Haidomyrmodes mammuthus*). In some cases, the length of flagellomeres diminishes towards the apex (*G. cretacica*) or slightly increases (*G. orientalis, G. occidentalis*). In *Sphecomyrma mesaki*, *Myanmyrma gracilis*, *Haidomyrmex zigrasi*, *Haidoterminus cippus*, and *G. subcuspis* it diminishes, then increases; in *Haidomyrmex scimitarus* it diminishes, then increases, and finally diminishes again. Thus, *G. cretacica* is unique in having flagellomeres which diminish in length towards the apex; this pattern resembles that of the Cretaceous males and *Paraponera*. *Haidomyrmex scimitarus* and *Haidomyrmodes mammuthus* also exhibit unusual antennae with a short terminal antennomere.

(3) Crown-group ant males: Antennae are filiform with similar segments, although in many cases the terminal segment is about 1.5 times longer than the others. In some cases, flagellomeres slightly increase in width and length towards the apex; in two cases (*Leptogenys* Roger, *Platythyrea* Roger), they diminish in length from F2/F3 towards the apex (the terminal flagellomere is again noticeably longer). Finally, in some genera (e.g., *Myrmica* Latreille) terminal flagellomeres are long and wide, so antenna is clavate.

(4) Cretaceous ant males: The length and probably width of flagelomeres diminish towards the apex; the terminal flagellomere is not longer than the penultimate one.

#### A comparison of the relative length of antennal parts

(1) The smallest scape length (relative to flagellum length) is found in the Cretaceous ant males, followed by the crown-group ant males, next the females of the Sphecomyrmini, Haidomyrmecini, and finally by the crown-group ant females. The relative scape length of the crown-group ant females is more than three times greater than that of the Sphecomyrmini, and two times greater than that of the Haidomyrmecini. Bolton (2003) provided a SL-to-FL ratio of 25% for the Sphecomyrminae, but the analysis conducted here shows the ratio of about 20% (Table S9).

(2) The pedicel of the crown-group ant females are approximately 20% longer than F1 and F2 (Table S16), but there are exceptions in which PL < F1L and PL = F1L (Table S2). F1L and F2L are equal statistically, but exceptions include F1L > F2L and F1L < F2L.

(3) The stem-group ant females, except for the Haidomyrmecini, and all males (both crown-group and Cretaceous) demonstrate a statistically significant pattern F1L > PL < F2L (Table S16). Moreover, the difference in length is often considerable: PL < F1L three-seven times in the Cretaceous ant males, up to seven times in the crown-group ant males, two-three times in the Sphecomyrmini, *Gerontoformica cretacica*, *Myanmyrma*; PL < F2L three-four times in the Cretaceous ant males, up to six times in the crown-group ant males, 1.5 times in *G. cretacica* and *Myanmyrma*, up to almost three times in the females of the Sphecomyrmini. The females show only three exceptions to the aforementioned pattern: PL = F2L (*Boltonimecia canadensis*, *Zigrasimecia ferox* #1), PL > F2L (*G. occidentalis*). More exceptions, including patterns PL > F1L, PL > F2L, PL = F2L, are seen in the males (Table S3). The comparison of F1L and F2L shows that only in the Sphecomyrmini F1L > F2L (without exception, and this pattern is confirmed statistically). In all other groups, F1L is not statistically different from F2L (Table S16). The greatest difference between F1L and F2L (up to two times) is found in the Sphecomyrmini, *G. cretacica*, *Myanmyrma, Archaeopone taylori*, *Baikuris mandibularis*, and some crown-group ant males.

(4) Unlike the Sphecomyrminae, but like the crown-group ant females, *Boltonimecia* demonstrates a pattern PL > F1L. However, unlike the crown-group ant females, *Boltonimecia* also demonstrates patterns PL = F2L and F1L < F2L (Table S2).

(5) The Haidomyrmecini is a heterogeneous group in terms of antennal metrics. They, unlike the crown-group ant females and Sphecomyrmini, did not show statistical differences between PL, F1L, F2L (Table S16), but show different patterns: in *Haidomyrmex scimitarus* and *H. cerberus* PL < F1L, in *Haidoterminus cippus* and *Haidomyrmodes mammuthus* PL = F1L, in *Haidomyrmex zigrasi* PL > F1L (all differences are minor, except for *Haidomyrmex scimitarus* showing a two-fold difference), in *Haidomyrmex zigrasi* and *Haidomyrmodes mammuthus* PL = F2L; in *Haidomyrmex scimitarus* and *H. cerberus* PL < F2L (a considerable difference of two and four times respectively), in *Haidoterminus cippus* PL > F2L, in *Haidomyrmex scimitarus* and *H. cerberus* F1L < F2L (1.5- and two-fold differences respectively) (*H. zigrasi* also shows F1L < F2L but with minor difference), in *Haidomyrmodes mammuthus* F1L = F2L, in *Haidoterminus cippus* F1L > F2L (minor difference) (Table S2).

#### A comparison of male and female indices

(1) Scape. The means of two indices (SL/HL, SL/FL) of the Cretaceous ant males are lower than those of the females of Sphecomyrminae, while the means of the crown-group ant males are almost equal to those of the females of Sphecomyrminae. Rare exceptions include the crown-group ant males and Cretaceous ant males with large indices, which are as large as in the crown-group ant females (Tables S2, S3, S4, S9, S15).

(2) Flagellum. The means of FL/HL of the Cretaceous ant males and crown-group ant males are greater than those of the stem- and crown-group ant females (Tables S5, S15).

(3) Pedicel. All pedicellar indices of the males studied are not different from those of the females (Tables S6, S10, S15).

(4) The first and second segment of the flagellum. The patterns of these segments resemble the scape patterns: the indices of the crown-group ant males are similar to those of the females of Sphecomyrminae, and noticeably greater than those of the crown-group ant females. The largest indices are seen in the Cretaceous ant males (Tables S7, S8, S11, S12, S15).

(5) The correlation between HL and length of different antennal parts was weaker in the males (R^2^ = 0.3-0.6), than in the females (R^2^ = 0.5-0.8). The correlation between SL and FL was negligible in the males (R^2^ = 0.02).

#### General observations on the shape of the pedicel

A long pedicel must be narrowed towards the scape and bent at its base to facilitate the scape-pedicel-flagellum articulation. Almost all crown-group ant females have the pedicels of such a shape, even those whose pedicels are not longer than F2 and F3. Rare exceptions include some army ants: their pedicels are short, with an almost non-existent bent, but are triangular (i.e., with narrowed bases) as in all other ants. Likewise, in some specimens of *Myrmecia* and *Nothomyrmecia* Clark the bent is almost non-existent, but the pedicel is always narrowed at the base. Crown-group ant males also have the pedicels mainly narrowed towards the scape and bent at the base, but the males with a short scape (SL/HL < 0.2) have barrel-shaped or spherical pedicels. Cretaceous males seem to demonstrate the same pattern: the only species with the scape index below 0.2 (*Baikuris mandibularis*) has a barrel-shaped pedicel, but all other Cretaceous males have triangular pedicels. All stem-group ant females have the scape index equal to or greater than 0.2 and so triangular pedicels. Thus, Dlussky and Fedoseeva’s (1988) assumption about the unique shape of pedicels in Sphecomyrminae is erroneous.

#### A comparison of the antennal indices of female ants and female non-ant Aculeata

(1) SL/AL of the crown-group ants is greater than that of all Vespoidea and Apoidea listed in Dlussky and Fedoseeva (1988) (i.e., crown-group ants have the greatest relative scape length) (Fig. 5A). The second greatest mean is seen in the social Vespoidea and Apoidea. Interestingly, the lowest SL/AL of the crown-group ants (in the Pseudomyrmicinae, Dorylinae, Leptanillinae, Martialinae) is almost equal to the lowest SL/AL of the social non-ant Hymenoptera (Table S2). The stem-group ants have one of the lowest mean of this index, and the lowest absolute value. The ANOVA indicated that the means were significantly different (F_4,109_ = 62.45, P < 0.0001). The planned comparisons showed that the differences between the crown-group ants and social Aculeata as well as between the stem-group ants and Vespoidea, Apoidea were negligible (Table S13). Thus, crown-group ants and other social Aculeata indeed underwent scape elongation (probably with simultaneous shortening of the flagellum), as was hypothesized by Dlussky (1983). These findings are the first statistical support in favor of Dlussky’s hypothesis.

(2) The stem-group ants have the greatest AL/HL mean (Fig. 5B), and also the two greatest absolute values of this index (Table S2). The ANOVA showed that the means were significantly different (F_4,109_ = 21.27; P < 0.0001), with statistically significant differences between all the groups, except for the difference between the crown-group ants and Vespoidea (Table S14).

**Figure 5.**
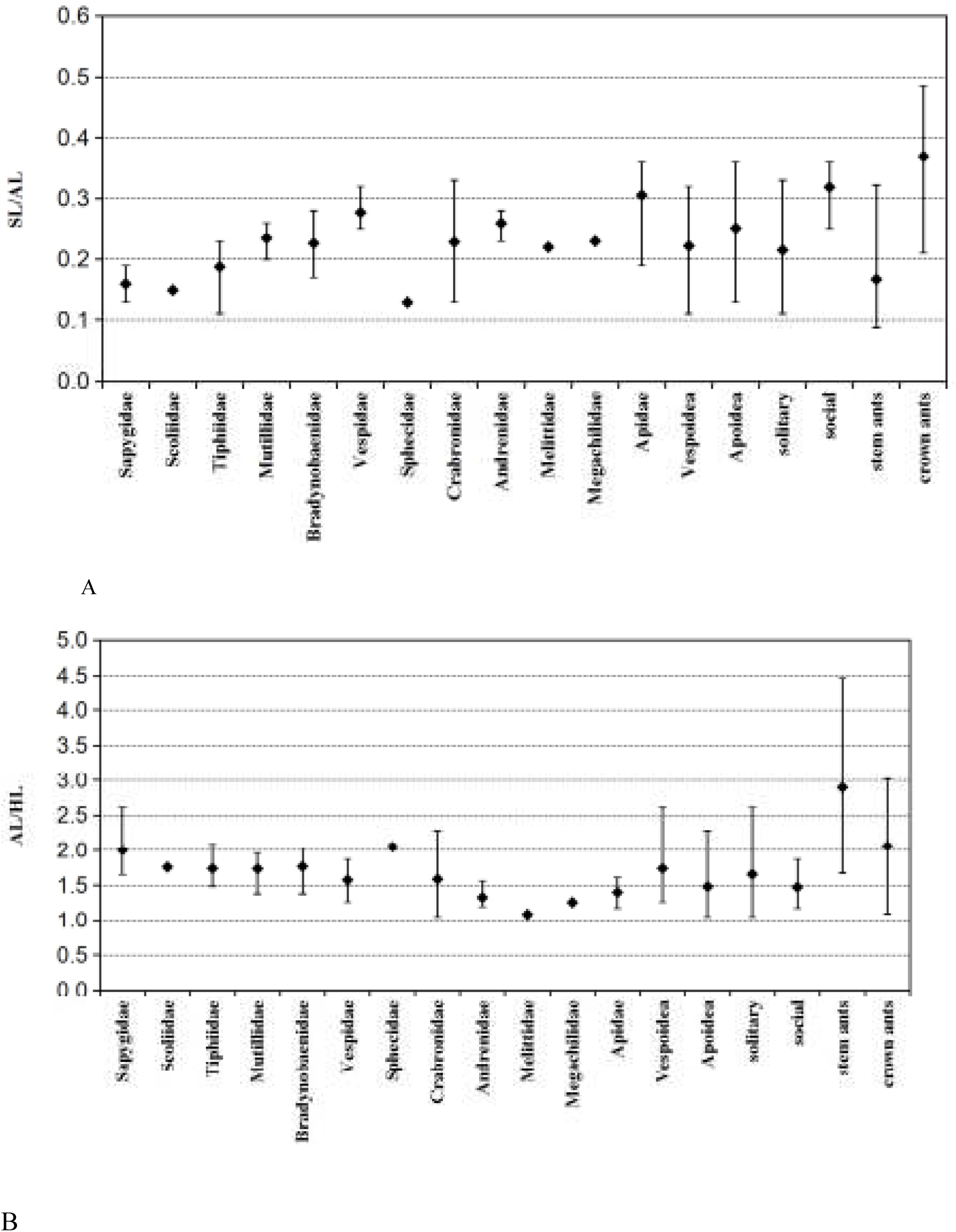
Range for the indices SL/AL and AL/HL in females of Vespoidea and Apoidea. **A)** Range for SL/AL. **B)** Range for AL/HL. (Note: Diamonds are arithmetic means. Data on Vespoidea and Apoidea are from Dlussky and Fedoseeva (1988), sorted by families according to the current classification. Vespoidea: families Sapygidae, Scoliidae, Tiphiidae, Mutillidae, Bradynobaenidae,Vespidae. Apoidea: families Sphecidae, Crabronidae, Andrenidae, Melittidae, Megachilidae, Apidae. Social Hymenoptera: *Vespa sp., Vespula sp., Polistes sp., Apis sp*. (1 species each), *Bombus sp.* (4 species). Stem ants: all stem-group ants from Table S1. Crown ants: all crown-group ants from Table S1).

#### Canonical discriminant analysis

This analysis based on the female indices SL/HL, FL/HL, F1L/HL, and F2L/HL was performed in order to achieve discrimination between crown- and stem-group ants. PL/HL was not used because it makes no contribution to the discrimination between the two groups; other unused indices are dependent on the aforementioned ones.

The first analysis was run with all the species. This approach explained 62% of the variation in the grouping variable (i.e., 62% of the species were correctly classified either as stem- or crown-group ants); on the other hand, the cross validated classification showed that 93% of the species were correctly classified (100% of crown- and 72% of stem-group ants). The discriminant function equation was as follows:

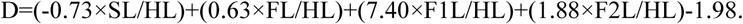

The mean discriminant scores (centroids): −0.90 for crown-group ants, 1.90 for stem-group ants (Fig. 6A; Table S17). The cut-off score separating both groups was 0. The discriminant function equation can be used to predict the group membership of newly discovered species by comparing the cut-off score, D and centroids. For example, three species described recently (Barden and Grimaldi 2016; Perrichot et al. 2016) have D scores 6.86 (*Gerontoformica maraudera* Barden and Grimaldi), 0.66 (*Camelomecia janovitzi*), and 2.75 (*Ceratomyrmex ellenbergeri* Perrichot, Wang and Engel) that classifies them as stem-group ants.

**Figure 6.**
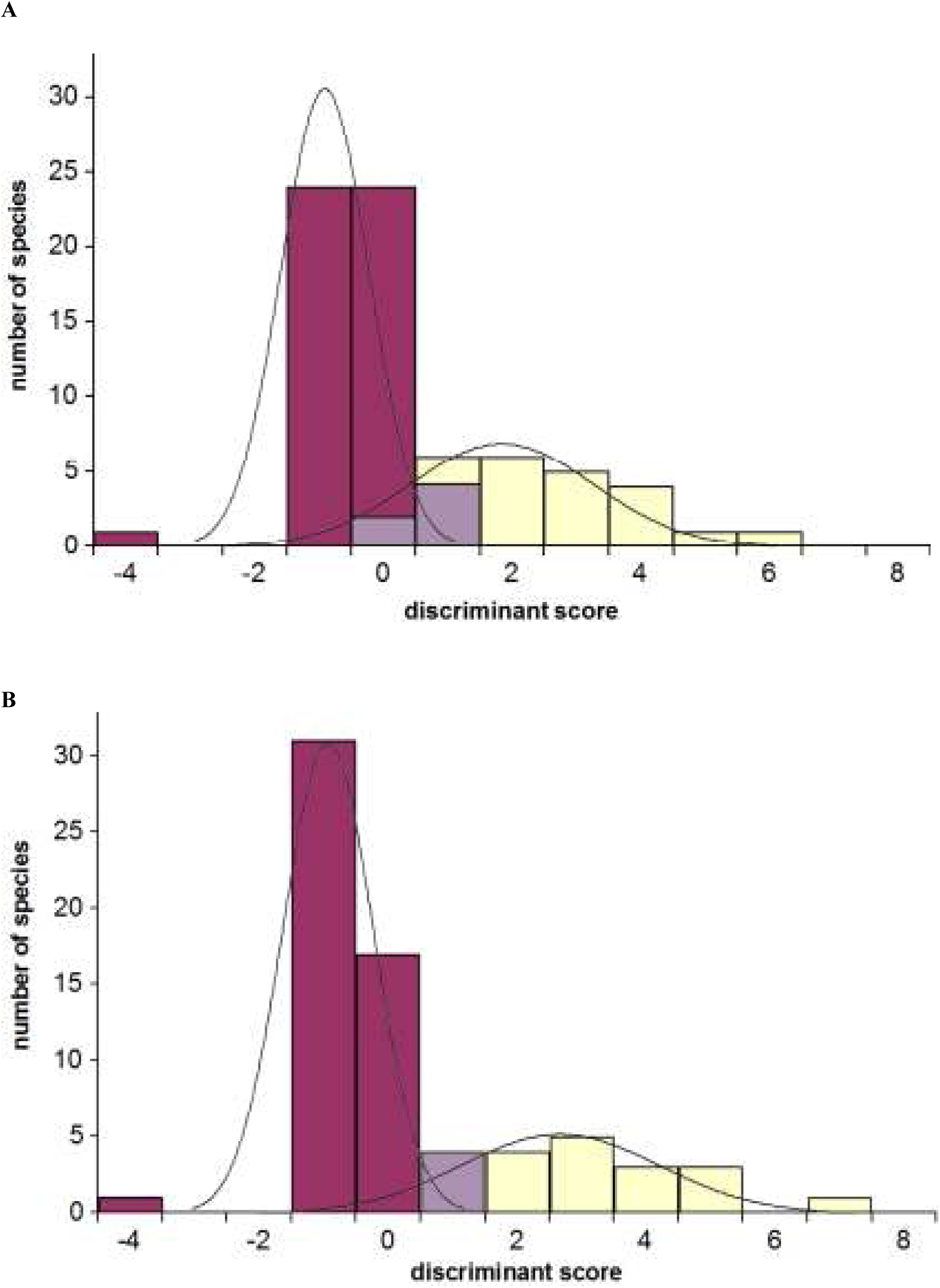
Results of canonical discriminant analysis. **A)** All species. **B)** Haidomyrmecini removed. (Note: Red violet bars are crown-group ants; yellow bars are stem-group ants; violet bars - area of overlap. Lines are Gaussian curves fitted to the data)

A better discrimination between the two groups can be achieved after the removal of the Haidomyrmecini because their discriminant scores overlap with the scores of the crown-group ants. This approach explained 72% of the variation and correctly classified 95% of the species (98% of crown-, 80% of stem-group ants) (Fig. 6B; Table S17). The discriminant function equation was as follows:

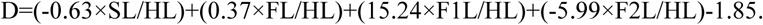

The centroids: −0.98 for crown-, 2.6 for stem-group ants. The cut-off score was 0.

Summing up these findings, antennal morphometry seems to be a promising heuristic tool in the taxonomy of fossil ants, as there exists a statistical basis for discriminating between stem- and crown-group ant females. The present research, however, should be viewed as preliminary, providing a basis for future studies when more stem-group species are discovered. New morphological data can be used in the model developed herein to confirm and extend its application.

### The higher classification of the ants

There is little doubt that the higher classification of the Formicidae with well-diagnosed stem and crown taxa will be an important development towards an understanding of the origin and evolution of the ants (Ward 2007). But here a number of complicating issues arise. First of all, we have to ask what ranks should be given to stem taxa, how we define the highest ranks of the Formicidae (in other words, what makes a subfamily a subfamily?), and finally, is it possible at all to apply the same classification principles to such different parts of the phylogenetic tree as crown and stem branches?

Obviously, even in modern groups, it is a challenge to find strong morphological characters of a subfamilial level. For example, a gastral constriction has been for a long time considered as such a character - it is one of the main characteristics of the Ponerinae in the old sense, or several subfamilies of the poneroids in the modern sense; but a gastral constriction is also present in the ectaheteromorphs, which have been assigned to the formicoid clade only after extensive molecular analyses (Moreau et al. 2006; Brady et al. 2006; Rabeling et al. 2008). The next example is a one- or two-segmented waist, which is a reliable subfamily-level character in some cases but variable in others (in the Dorylinae, Leptanillinae, Myrmeciinae).

Not surprisingly, the situation in stem-group ants is much more complicated, because the “weight” of a character decreases the farther we move down the evolutionary scale. For instance, it is possible that the presence of a gastral constriction is a genus-level character in the Sphecomyrminae, unlike crown-group ants, where it varies at a subfamilial level (discussed below). As Dlussky and Fedoseeva (1988) noted, the ancestors of all branches of the ant tree were probably so similar that, if they existed today, we would have assigned all of them to a single genus. This means that in ancestral groups, “strong” morphological characters could be not only vague but also combined in unusual ways.

Probably realizing that morphology is a poor tool for defining a taxon’s rank, Hennig (1966) suggested that ranks could be associated with the absolute age of the taxa. If we check this assumption by the time needed to accumulate enough distinguishing characters, the following points can be outlined.

The youngest subfamilies of the crown clade are at least 70 Ma old. For example, the dorylomorph clade represented by six subclades, which, until recently, had the status of subfamilies, but now are subsumed into a single subfamily (Dorylinae), is 74-101 Ma old (Brady et al. 2014). It is revealing that it took several decades of morphological studies to justify the split of the dorylomorphs into six subfamilies (Baroni Urbani et al. 1992; Bolton 2003; Brady and Ward 2005), and the first molecular data confirmed the split (Brady et al. 2006; Moreau et al. 2006; Rabeling et al. 2008); but deeper molecular research showed that this conclusion was erroneous (Brady et al. 2014). It seems plausible that an age greater than 70 Ma is required for a group to be confidently recognized as an ant subfamily.

From this, only if the age of the Sphecomyrminae is about 150 Ma, and if the Sphecomyrminae survived until the end of the Cretaceous, their subfamily rank can be congruent with modern subfamilies (not taking into account the fact that the rate of evolutionary radiation of Cenozoic ants (i.e., crown-group ants) was higher than that of Mesozoic ants (mainly stem-group lineages) and thus the Sphecomyrminae probably required even more time to accumulate characters of a subfamilial “weight” than crown-group ants). But as inferred from fossil records, the Sphecomyrminae existed for only 20 Ma.

The inability to apply the same criteria for the classification of stem and crown groups raises the problem of a classification system. Indeed, in the Linnaean system, it is usually impossible to classify stem taxa, as only the taxa that are explicitly associated with ranks can be formally named (Ereshefsky 2001; Pleijel and Rouse 2003; Joyce et al. 2004). To give ants as an example, under the Linnaean system, stem clades and supra-subfamilial clades of the crown clade (e.g., formicoids, poneroids) can be named only if assigned to intermediate ranks, which will make the classification exceedingly complicated.

Another problem is that the rank-based system implies comparability across taxa of the same rank, but in fact these taxa lack equivalency (Ereshevsky 2001), they are not comparable entities in a cladistic sense (Pleijel and Rouse 2003; Dubois 2007). If so, the ranks themselves are subjective devices, serving only to build classification; they are arbitrary and lack biological meaning (Hennig 1966).

The alternative of the Linnaean system is a rankless system such as the one governed by the PhyloCode (Cantino and de Queiroz 2010). Phylogenetic nomenclature of the PhyloCode extends the concept of tree thinking to biological nomenclature (Baum and Smith 2012) and, by classifying all organisms, not just descendants, helps, among other things, to explain the relationships between stem and crown branches.

To continue the example given above, rankless classification helps to clarify the status of the Sphecomyrminae. Their taxonomic status is indeed an unsolvable dilemma in the Linnean system: although evidence of the long (at least 70 Ma) evolution of the Sphecomyrminae is absent, their phylogenetic level is expected to be higher than that of any modern subfamily. Since a stem taxon of a higher taxon is identical to that higher taxon (Hennig 1966), the Shecomyrminae and other stem taxa can be viewed as sister groups to the crown clade; they are not comparable to the subfamilies of the crown clade but are equivalent to the crown clade as a whole.

Similarly, a phylogeny-based classification allows to place the Cretaceous representatives of crown-group ants, for which using Linnaean ranks is also problematic, in particular clades. The statistical analysis of the antennal indices shows that *Kyromyrma neffi*, *Canapone dentata*, *Eotapinoma macalpini*, *Chronomyrmex medicinehatensis*, and *Brownimecia clavata* Grimaldi, Agosti and Carpenter undoubtedly belong to crown-group ants, but these first crown-group ants cannot be placed in any extant tribe; they are most probably stem taxa of extant subfamilies.

In a proposed higher classification of the ants, by using phylogenetic and Linnaean nomenclature together, I attempt to show how clades that are important for our understanding of ant evolution but are not explicitly associated with Linnaean ranks can be formally defined. I believe that the two systems should not be viewed as competitors, but instead as complementing each other.

### Crown-Formicidae

#### Diagnosis (workers, gynes)

(1) Scape long (SL/HL no less than 0.3, often more than 0.6; SL/FL more than 0.3, often more than 0.5); (2) flagellomeres short (F1L/HL and F2L/HL less than 0.20, often less than 0.10) and whole flagellum short (FL/HL often < 1.4); (3) terminal flagellomere usually elongated; (4) PL often > F1L and > F2L (20% on average); (5) F1L = F2L, middle flafellomeres often equal in length as well; (6) antenna clavate or at least flagellomeres gradually widening to apex; (7) mandibles broad, multidentate; (8) clypeus posteriorly inserted between antennal sockets or not; (9) two spurs of mesotibia and metatibia present or not; (10) preapical tooth on pretarsal claws present or not; (11) trochantellus absent; (12) waist one or two-segmented.

#### Comment

Under the PhyloCode, the taxon should be named “*Formicidae*^P^ Stephens 1829, converted clade name” and defined as “the clade originating with the most recent common ancestor of *Martialis heureka* Rabeling and Verhaagh, 2008 and *Formica rufa* Linnaeus, 1761”. This clade corresponds to the family Formicidae Latreille, 1809, excluding stem taxa.

#### Composition

The clade is monophyletic (Baroni Urbani et al. 1992; Brady et al. 2006), with estimates of diversity of more than 13 000 species. The high-level classification of the Formicidae has been greatly revised by Bolton (2003) and then received further support from molecular phylogenies (Brady et al. 2006; Moreau et al. 2006; Rabeling et al. 2008). The composition of the subclades that correspond to 17 monophyletic subfamilies is stable (Ward 2011), but the clade structure above the subfamilial level is not well understood. It is believed that the clade splits into two major subclades: the formicoid clade (formicoids) and the poneroid clade (poneroids) (Ward 2007). The former is a stable clade, although its structure is different in morphological and molecular analyses; the latter is either a sister clade to the formicoids or a paraphyletic group from which the formicoids arose (Ward 2011).

#### Discussion

*Formicidae*^P^ have antennae with three functional parts: a long scape, short middle part, elongated/thickened apical part. The scape comprises about 60% of the length of the flagellum (three times greater than in stem-group ants); the flagellum is about 1.3 times longer than the head (1.8 times in stem-group ants).

The first *Formicidae*^P^ are known from the Cretaceous: *Kyromyrma neffi* and *Brownimecia clavata* are from New Jersey amber (Turonian, 92 Ma); *Eotapinoma macalpini, Chronomyrmex medicinehatensis,* and *Canapone dentata* are from Canadian amber (Campanian, 78–79 Ma).

*B. clavata* almost certainly belongs to *Formicidae*^P^, but its position within the clade remains unclear. It may be a stem taxon of either the poneromorphs or the poneromorphs + leptanillomorphs (Bolton 2003), or even a stem taxon of a subfamily (such as the Amblyoponinae or Ponerinae). This species is not included in the statistical analysis due to its uncertain taxonomic position; however *Brownimecia*’s indices are close to the means for the crown-group ants, which at least confirms its affiliation with the crown clade.

The taxa for which placement (within or outside of *Formicidae*^P^) is highly uncertain: *Cananeuretus occidentalis, Burmomyrma rossi* Dlussky, Formiciinae. *C. occidentalis’* (Aneuretinae) index SL/HL is close to the mean for the crown group, but the index PL/HL is more than twice greater than the greatest values in the stem- and crown-group ants, and more than four times greater than their means (Fig. 4D). It is not possible to calculate the other indices due to the incomplete preservation of the antennae. *B. rossi* is thought to be a stem group of the Aneuretinae (Dlussky 1996), but poor preservation makes this hard to confirm. The early Eocene (50 Ma) (Archibald et al. 2011) extinct subfamily Formiciinae was placed outside of the crown-group ants in the cladistic study of Baroni-Urbani et al. (1992) and Grimaldi et al. (1997). Those data should be interpreted with caution, because the Formiciinae are known only from poorly preserved imprints in rock. Their antennae are not preserved, so the Formiciinae were not included in the statistical analysis performed here.

### Pan-Formicidae

#### Diagnosis (workers, gynes)

Characters shared by crown-group ants (*Formicidae*^P^) and stem-group ants: (1) wingless worker caste; (2) head prognathous; (3) metapleural gland present; (4) differentiated petiole present. Characters of crown-group ants (*Formicidae*^P^): see Crown-Formicidae. Characters of stem-group ants: (1) scape short (SL/HL can be less than 0.3, rarely more than 0.6; SL/FL often less than 0.3 and never more than 0.5); (2) flagellomeres long (F1L/HL and F2L/HL > 0.10, often more than 0.20) and whole flagellum long (FL/HL > 1.4); (3) terminal flagellomere elongated or not; (4) often F1L > PL < F2L; (5) F1 longer, shorter, or equal in length to F2; (6) antenna without club, flagellomeres rarely widened/elongated towards apex, may diminish in length towards apex; (7) mandibles narrow, linear bidentate or highly specialized (L-, scythe-shaped, cuplike) monodentate; (8) clypeus posteriorly usually not inserted between antennal sockets; (8) trochantellus present or not; (9) mesotibia and metatibia each with two spurs; (10) preapical tooth on pretarsal claws often present; (11) waist one-segmented.

#### Comment

Under the PhyloCode, the taxon should be named “*Pan-Formicidae*^P^, new clade name” and defined as “the total clade composed of the crown clade *Formicidae*^P^ and all extinct species that share a more recent common ancestor with *Formicidae*^P^ than with any extant species that are not members of *Formicidae*^P^”. Reference phylogenies: Grimaldi et al. 1997: Barden and Grimaldi 2016. More characters of crown-group ants are listed elsewhere (Bolton 2003; Boudinot 2015).

#### Composition

*Formicidae*^P^*, Sphecomyrminae*^P^.

#### Discussion

In stem-group ants, unlike crown-group ants, the antenna is not divided into three functional parts; usually there is no striking difference between the length of the scape and flagellomeres; the antennal club is absent.

The taxa (genera) which definitely belong to *Pan-Formicidae*^P^ but whose position within the clade remains unclear: *Archaeopone* Dlussky, *Baikuris* Dlussky, *Dlusskyidris* Bolton, *Poneropterus* Dlussky, *Myanmyrma* Engel and Grimaldi, *Camelomecia* Barden and Grimaldi. The first four genera are known from males only. *Myanmyrma* (as well as some species of *Gerontoformica* - see below) may represent a separate clade of *Pan-Formicidae*^P^*. Myanmyrma*’s indices are strongly different from those of the crown group. SL/HL is close to that of the Sphecomyrmini, but SL/AL is the lowest of all Aculeata species considered in the present study (i.e., *Myanmyra* has one of the shortest scapes). Regarding other antennal parts, *Myanmyrma* is also unique: it has the longest pedicel, F1, F2, and one of the longest flagella. Other unique characters include extremely elongated mandibles, a bilobate clypeal margin, and long genal process (all are putative apomorphies). The “poneroid” habitus of *Myanmyrma* (Engel and Grimaldi 2005) is questionable; its deep gastral constriction, implying a morphologically differentiated postpetiole (unknown in stem-group ants), may be an artifact. Thus, it is safe to discard the assumption that *Myanmyrma* belongs to crown-group ants (Engel and Grimaldi 2005). All the evidences indicate that *Myanmyrma* is a stem-group species, showing a unique combination of characters. The same may apply to the recently described *Camelomecia* (Barden and Grimaldi 2016).

### Subfamily Sphecomyrminae Wilson and Brown 1967

#### Composition

Tribes Sphecomyrmini, Boltonimeciini, Haidomyrmecini.

#### Discussion

It has been assumed that the Sphecomyrminae is a group close to the formicoid clade of the crown group (Taylor 1978; Rabelling et al. 2008). The Sphecomyrminae do indeed resemble representatives of the formicoid clade by the habitus, the presence of ocelli, and an unspecialized (though only in the Sphecomyrmini) head capsule similar to that of primitive formicines (*Prolasius* Forel, *Notoncus* Emery, *Prenolepis* Mayr) (Wilson et al. 1967). However, the data presented here suggest that the Sphecomyrminae is a stem clade, i.e., a sister group to the crown clade.

The clade is doubtfully monophyletic. The three subclades (tribes) may represent an artificial assemblage, although the Sphecomyrmini and Boltonimeciini do seem closely related. Some important characters appeared to be variable. For example, scape length in Sphecomyrminae comprises 20% of flagellum length, compared with 60% in the crown group, and varies in a large range (discussed below). Other variable characters include the presence of the gastral constriction, trochantellus, and clypeal peg-like setae.

### Tribe Sphecomyrmini Wilson and Brown, 1967

#### Diagnosis (workers, gynes)

(1) head capsule unspecialized; (2) mandibles unspecialized; (3) anterolateral clypeal margins not produced over mandibular bases in rounded lobes; (4) peg-like setae on anterior clypeal margin present or not; (5) ocelli present; (6) F1L > PL < F2L, F1L > F2L; F1 often longest flagellomere; (7) neck short or long; (8) petiole subsessile or pedunculate; (9) gastral constriction present or not.

#### Comment

Under the PhyloCode, the taxon should be named “*Sphecomyrmini*^P^, converted clade name” and defined as “the clade consisting of *Sphecomyrma freyi* Wilson and Brown, 1967 and all species that share a more recent common ancestor with *Sphecomyrma freyi* Wilson and Brown, 1967 than with *Haidomyrmex cerberus* Dlussky, 1996 or *Boltonimecia canadensis* (Wilson 1985)”.

#### Composition

Genera *Sphecomyrma* Wilson and Brown (type genus), 1967, *Cretomyrma* Dlussky, 1975, *Armania* Dlussky, 1983, *Pseudarmania* Dlussky, 1983, *Orapia* Dlussky, Brothers and Rasnitsyn, 2004, *Gerontoformica* Nel and Perrault, 2004.

#### Dicussion

The species of this tribe seem to be less specialized than those of the other two tribes of the Sphecomyrminae. Also, the tribe is heterogeneous due to the presence of possibly non-monophyletic genera *Sphecomyrma* and *Gerontoformica*.

In the genus *Sphecomyrma*, the SL/HL index has an unusually large range. Even in a single species, *S. freyi,* values range from 0.28 to 0.62 (Table S2), i.e., the difference is 120%. Data on 11 modern genera (Radchenko 1991, 1994; Seifert 1992, 2000, 2003; MacKay 1993; Radchenko and Elmes 1998, 2003; Ward 1999; Radchenko et al. 2002; Baroni Urbani and de Andrade 2003; Wild 2004; Bolton 2007; Bolton and Fisher 2011) show that the range of the SL/HL among workers, gynes, and workers + gynes within a given species rarely exceeds 10%, with a single maximum of 20% in *Myrmica* (Radchenko 1994), 31% in *Linepithema* Mayr (Wild 2004), 31% in *Dolichoderus* Lund (MacKay 1993). Such results raise the possibility that *S. freyi* #3, which has the SL/HL value of 0.28, does not belong to *Sphecomyrma*.

Regarding the variation of the SL/HL index within a genus, the findings are the following. In the Cretaceous ant males as exemplified by *Baikuris,* the range is 77%; in the workers of *Sphecomyrma,* it is 160% (or 106% if the clypeal lobe of *S. mesaki* is excluded). In modern genera, males are characterized by a large range: 100% in *Proceratium* Roger (Baroni Urbani and de Andrade 2003), 204% in *Dolichoderus* (MacKay 1993), 175% (Seifert 1988) and 260% (Radchenko 1994) in *Myrmica.* The workers and gynes of modern genera have range from 30% in *Pseudomyrmex* Lund to 55% in *Proceratium* (Radchenko 1994; Seifert 1992, 2000, 2003; Baroni Urbani and de Andrade 2003; Bolton and Fisher 2011; Ward 1999); only in the Dolichoderinae there are single large values, causing larger ranges up to 128% in *Dolichoderus* (MacKay 1993), and to 121% in *Technomyrmex* Mayr (Bolton 2007). Therefore, whereas the range in *Baikuris* is consistent with the data on recent genera, the range in *Sphecomyrma* is not. Although the limits of within-genus variation are not known for Cretaceous representatives, taking into account the unique morphology of *S. mesaki* (short scape, antennal scrobes), its taxonomic position should also be re-examined.

The genus *Gerontoformica*, which includes 13 species, is even more problematic. Indeed, *Gerontoformica* is variable in generic- and higher-level characters, such as the scape length, palp formula, the presence of a petiolar peduncle, ocelli, gastral constriction, and trochantellus. In some species the head seems to be specialized as in the Boltonimeciini (frontal part thickened, anterolateral clypeal lobes present). All the species have peg-like setae on the anterior clypeal margin, again as in the Boltonimeciini. The SL/HL index ranges from 0.19 (the minimum value obtained in this study) to 0.67 (one of the highest values of all stem-group ants) (Table S2), i.e., the difference is 250%, which is larger than any within-genus range obtained here.

Recently (Barden and Grimaldi 2016) *Gerontoformica* and *Sphecomyrmodes* Engel and Grimaldi have been synonymized, which made the genus even more heterogeneous. Of special interest is the species *G. cretacica* transferred from incertae sedis. In terms of the results of the statistical analysis, *G. cretacica* occupies an intermediate position between the crown- and stem-group ants. For SL/HL, *G. cretacica* is similar to the crown-group ants; its scape is the longest of all stem-group ants known, but its flagellum is also elongated proportionally to the scape, and thus the indices FL/HL and AL/HL are among the largest reported. The same applies to the pedicel, F1 and F2. Other interesting features are: the terminal flagellomere is not elongated (a rare character also found in two species of the Haidomyrmecini), antenna without the club, flagellomeres diminish in length towards the apex (Nel et al. 2004).

It is possible that *Gerontoformica* can be split into several genera, and these genera can be placed in different tribes within the Sphecomyrminae. But *G. cretacica* may fall out of *Sphecomyrminae*^P^; its unique combination of characters (if it is not an artifact of preservation as suggested by the latest report - Barden and Grimaldi 2016) may indicate that it is a representative of a subclade of *Pan-Formicidae*^P^ branched close to the crown clade.

The problem of large within-genus variations of the scape length seen in *Sphecomyrma* and *Gerontoformica* may have, however, another angle. There is a long-lasting debate on whether stem-group ants were eusocial or not, and, as it was already pointed out, the answer to this question is probably linked to the problem of scape elongation (Dlussky 1983; Dlussky and Fedoseeva 1988). It is reasonable to assume that the transition to eusociality did not occur instantly but gradually, via the stage of facultative sociality, i.e., social behavior in stem-group ants depended on, for example, abiotic environmental conditions and varied even among closely related species. A similar pattern of social organization is found in the modern halictine bees (Yanega 1997). If this is the case, then the scape length was not stable in stem lineages.

Lastly, I will review the taxonomic status of the Armaniinae, an enigmatic group known mainly from winged forms preserved as imprints in rock. Dlussky initially assigned all his new Cretaceous species to the Sphecomyrminae (Dlussky 1975) but then transferred to the new family Armaniidae (Dlussky 1983). Wilson returned them to the Sphecomyrminae, and synonimized almost all genera of the Armaniinae (except *Cretomyrma*) with the genus *Sphecomyrma* (Wilson 1987). Bolton changed the status of Dlussky’s family to the subfamily Armaniinae (Bolton 1994, 2003); Dlussky first accepted Bolton’s approach (Dlussky 1996) but then again called the group Armaniidae (Dlussky 1999b), and then again mentioned the subfamily Armaniinae (Dlussky *et al* 2004). In 2005, Wilson mentioned this group as the family Armaniidae (Wilson and Hölldobler 2005). Such confusion is caused by the absence of reliable characters distinguishing the Armaniinae from the other groups, especially from the Sphecomyrminae.

The gastral constriction is one of the most confusing characters. For example, *Dolichomyrma* was described without (Dlussky 1975) or with (Wilson 1987) the gastral constriction; *Armania* - with (Dlussky 1983) or without (Wilson 1987); *Arhaeopone* - with (Dlussky 1975) or without (Dlussky 1983); *Petropone* most probably has the gastral constriction (Dlussky 1975). It is now clear, however, that the gastral constriction is not a stable character in Sphecomyrminae, being present in some genera and absent in others. The trochantellus may be an important character distinguishing the Armaniinae and Sphecomyrminae (Dlussky 1983), but it not stable either, being variable even within a genus (e.g., in *Gerontoformica*) (Barden and Grimaldi 2014).

Also, the Armaniinae are characterized by a “very short scape”, in contrast to just a “short scape” of the Sphecomyrminae (Bolton 2003). This is viewed as one of the most important diagnostic characters of the Armaniinae (Dlussky 1983). However, as can be seen from the statistical analysis, the scape indices as well as other antennal indices of the Armaniinae are not at all different from those of the Sphecomyrminae or Sphecomyrmini (Tables S2, S4-S8; Fig. 4). To give more support to these findings, I compared within-subfamily ranges of the scape index SL/HL in workers. The difference between the extreme values was the following: 3.5 times in the Sphecomyrminae (between *Gerontoformica orientalis* and *G. rugosus*); 2.2 times in the Ponerinae (from 0.53 in *Feroponera* Bolton and Fisher to 1.16 in *Diacamma* Mayr) (Shattuck and Barnett 2006; Bolton and Fisher 2008); 3.4 times in the Dolichoderinae (from 0.65 in *Anillidris* Santschi to 2.3 in *Leptomyrmex*) (Shattuck 1992; Lucky and Ward 2010; Schmidt et al. 2013); 4.6 times in the Myrmicinae (from 0.28 in *Metapone* Forel to 1.3 in *Aphaenogaster* Mayr) (Alpert 2007; Shattuck 2008); 6.2 times in the Formicinae (from 0.34 in *Cladomyrma* Wheeler to 2.13 in *Euprenolepis* Emery) (Agosti et al. 1999; LaPolla 2009). The difference observed between the Armaniinae and Sphecomyrminae is much lower than these values (Table S4). Thus even in comparison with diverse modern groups, the Sphecomyrmini and Armaniinae can be viewed as one homogeneous entity.

Another important feature listed in the diagnosis of the Armaniinae is the shape of the petiole. In the Armaniinae, the petiole is broadly attached posteriorly to the gaster, which resembles the petiole of non-ant Vespoidea (e.g., Sierolomorphidae), while in the Sphecomyrminae it is nodiform (Dlussky 1999b). This may be the only remaining argument against changing the taxonomic status of the Armaniidae/Armaniinae, because such an attachment is unique and obviously plesiomorphic.

The drawings and photos of the Armaniinae, however, do not speak in favor of this conclusion. For example, in *Orapia*, *Pseudoarmania*, *Armania curiosa* the petiole looks rounded above and on the sides, not different from the petiole of the Sphecomyrminae. In the putative males of the Armaniinae (genera *Archaeopone* and *Poneropterus*), the petiole is massive but again similar to that of the putative males of the Sphecomyrminae (*Sphecomyrma* and *Dlusskyridris*). From the poorly preserved imprints of *Khetania*, *Armania pristina*, and *A. capitata*, one may conclude that the petiole does not appear to be differentiated from the gaster, which raises the question about the assignation of these specimens to the ants.

The only exception is a well-preserved imprint of *Armania robusta* with a relatively massive petiole, which seems, indeed, to be broadly attached to the gaster (although partly hidden by a coxa). Interestingly, Wilson (1987) considered this particular specimen as a gyne of *Sphecomyrma*, and its massive petiole as a sexually dimorphic feature, i.e., not as a subfamilial or even species-level character. Taking into account that there is no definition of a “broad attachment”, it is necessary to fill this gap by calculating the index PG/PH in the *A. robusta* holotype as well as in several species of stem- and crown-group ants. The values of this index in the ants with a nodiform petiole are the following: 0.5 (*Gerontoformica subcuspis*), 0.6 (*Sphecomyrma freyi*), 0.7 (*Boltonimecia* and gynes of *Leptanilla*). The larger value (indicating that the petiole is more broadly attached to the gaster) is found in *A. robusta*; but in the workers and gynes of *Stigmatomma* (Amblyoponinae) values vary from 0.7 to 0.9. It is therefore impossible to reach the unambiguous conclusion that the petiole of *A. robusta* is a unique feature in terms of the width of its attachment to the gaster.

In summary, it is hard to disagree with Wilson (1987) that the Armaniinae do not have any subfamilial-level feature, to say nothing of a familial one. So my interpretation of the taxonomic status of the Armaniinae is similar to that of Grimaldi et al. (1997). Relatively well-preserved genera used in the statistical analysis (*Armania*, *Pseudoarmania*, *Orapia*) were transferred to *Sphecomyrmini*^P^. All the other genera were considered to be groups of uncertain taxonomic position within *Formicidae*^P^ or Aculeata.

### Tribe Haidomyrmecini Bolton, 2003

#### Diagnosis (workers, gynes)

(1) head capsule specialized: face and genae concave, clypeus modified (putative apomorphy); (2) mandibles long, scythe or L-shaped (putative apomorphy); (3) anterolateral clypeal margins not produced over mandibular bases in rounded lobes; (4) peg-like setae on anterior clypeal margin absent; (5) ocelli present or not; (6) relative length of antennomeres variable; (7) terminal flagellomere elongated or not; (8) neck long; (9) petiole pedunculate; (10) gastral constriction present or not.

#### Comment

Under the PhyloCode, the taxon should be named “*Haidomyrmecini*^P^, converted clade name” and defined as “the clade consisting of *Haidomyrmex cerberus* Dlussky, 1996 and all species that share a more recent common ancestor with *Haidomyrmex cerberus* Dlussky, 1996 than with *Sphecomyrma freyi* Wilson and Brown, 1967 or *Boltonimecia canadensis* (Wilson 1985)”.

#### Composition

Genera *Haidomyrmex* Dlussky, 1996 (type genus), *Haidomyrmodes* Perrichot, Nel, Néraudeau, Lacau, Guyotet, 2008, *Haidoterminus* McKellar, Glasier and Engel, 2013, *Ceratomyrmex* Perrichot, Wang and Engel, 2016.

#### Discussion

This group is morphologically compact, and readily distinguishable from the other stem taxa. The unique head and mandibles are the most important morphological features, but some findings from the statistical analysis are also noteworthy. The scape indices as well as the index F1L/HL occupy an intermediate position between the Sphecomyrmini and crown-group ants. This negatively influenced the discrimination between stem- and crown-group ants in the canonical discriminant analysis, as well as limits the potential of antennal metrics for providing a more rigorous diagnosis of *Pan-Formicidae*^P^. All the other indices are statistically similar to those of the Sphecomyrmini. Unlike the Sphecomyrmini, there is no pattern F1L > P < F2L (many exceptions); however, some patterns in a F1-to-F2 ratio can be noted at the generic level: *Haidomyrmex* - F2L > F1L, *Haidomyrmodes* - F2L = F1L, *Haidoterminus* and *Ceratomyrmex* - F1L > F2L.

The presence of the gastral constriction in the Haidomyrmecini is questionable. *Haidomyrmodes* has been described with the gastral constriction, but it may be an artifact of preservation. It is also possible that this character is variable at the generic level, as in the Sphecomyrmini.

### Тribe Boltonimeciini trib.n

#### Diagnosis (workers)

(1) head capsule specialized: shield-like, with dorsal part thick and raised (putative apomorphy); (2) anterolateral clypeal margins produced over mandibular bases in rounded lobes (putative apomorphy); (3) peg-like setae on anterior clypeal margin present; (4) ocelli absent; (5) relative length of antennomeres variable; (6) neck long; (7) protibia with three spurs: one pectinate and two simple; (8) petiole pedunculate; (9) gastral constriction absent.

#### Comment

Under the PhyloCode, the taxon should be named “*Boltonimeciini*^P^, converted clade name” and defined as “the clade consisting of *Boltonimecia canadensis* (Wilson 1985) and all species that share a more recent common ancestor with *Boltonimecia canadensis* (Wilson 1985) than with *Sphecomyrma freyi* Wilson and Brown, 1967 or *Haidomyrmex cerberus* Dlussky, 1996”.

#### Composition

Genera *Boltonimecia* gen.n. (type genus), *Zigrasimecia* Barden and Grimaldi, 2013.

#### Discussion

A short scape, long flagellum, two spurs on meso- and metatibia, pretarsal claws with a preapical tooth, and the clypeus posteriorly not inserted between the antennal sockets indicate that *Boltonimecia* belongs to stem-group ants and the Sphecomyrminae. The only problem here is the form of the metapleural gland orifice and the presence of the mesoscutum and scutellum; these characters have been listed in the diagnosis of the Sphecomyrminae (Bolton 2003), but cannot be observed in *Boltonimecia* due to compression of the mesosoma.

The antennal indices of *Boltonimecia* and *Zigracimecia* (type species: *Z. tonsora*) are similar to those of the Sphecomyrmini, although some important peculiarities need to be mentioned. *Boltonimecia*’s SL/HL is one of the greatest among the Sphecomyrminae and is close to that of the crown-group ants. Both genera have a longer pedicel compared with the Sphecomyrmini; in *Boltonimecia*, the pedicel is so elongated that PL > F1L, as in the crown-group ants (Sphecomyrmini always have F1L > PL). In *Boltonimecia* and *Z. ferox* #1, PL = F2L (in Sphecomyrmini, PL < F2L, except for *Gerontoformica occidentalis*). Also, in *Boltonimecia*, F1L < F2L (in Sphecomyrmini, F1L > F2L without exception).

Therefore, the statistical analysis also leaves no doubt about the assignation of *Boltonimecia* and *Zigrasimecia* to stem-group ants and the Sphecomyrminae. These two genera, however, seem to be closer to each other than to the Sphecomyrmini, thus I propose that they be placed into a separate tribe.

The shield-like head, anterolateral clypeal margins produced over mandibular bases in rounded lobes, and three protibial spurs are the most important characters of the new tribe. Noteworthy, these are found also in some *Gerontoformica* species (Barden and Grimaldi 2014). In particular, the rounded anterolateral clypeal margins are present in *G. spiralis* and *G. tendir;* three protibial spurs (called “stiff setae” by the authors) - in *G. spiralis*, *G. subcuspis*, *G. magnus*, *G. rubustus*; at least some *Gerontoformica* have the shield-like head. Also, like Boltonimeciini, *G. rugosus* and *G. spiralis* have a long pedicel; all *Gerontoformica* have the peg-like setae on the anterior clypeal margin. These peculiarities are starting points for a thorough revision of a seemingly non-monophyletic genus *Gerontoformica*, which hopefully will be achieved in the near future.

The last part of a proposed higher classification is the subclade composition of the crown group *Formicidae*^P^. Since the composition of the subclades that correspond to monophyletic subfamilies is stable, a phylogeny-based classification is straightforward and congruent with the Linnaean system. In a formal phylogenetic definition of the subclades given in Table S18, attention was paid to the recommendation that the type species (article 11.7 of the PhyloCode) and the species used in the reference phylogenies (article 11.8 of the PhyloCode) should preferably be used as specifiers. For most of the clades, a node-based definition is used, except for the clades consisting of only one extant species, for which a branch-based definition is provided.

The vast majority of Cenozoic fossil crown-group ants fit nicely into the clades that correspond to tribes or genera in the Linnaean system; some Cretaceous crown-group species (*Brownimecia*) are most probably stem taxa to a clade of a higher (supra-subfamilial) taxonomic level; and only a few fossil crown-group species are related to modern subfamilies but do not fall into any of the tribes (i.e., they are stem taxa to those subfamilies). Most likely, only five taxa (genera) can be considered as forming pan-clades with three recent subfamilies, a formal definition of which is provided below (diagnoses after Bolton 2003, with modifications).

### Pan-Formicinae

#### Diagnosis (workers, gynes)

(1) acidopore present at apex of hypopygium (apomorphy); (2) sting absent; (3) helcium attached low on anterior face of first gastral segment.

#### Comment

Under the PhyloCode, the taxon should be named “*Pan-Formicinae*^P^, new clade name” and defined as “the total clade composed of the crown clade *Formicinae*^P^ and all extinct species that share a more recent common ancestor with *Formicinae*^P^ than with any extant species that are not members of *Formicinae*^P^”.

#### Composition

*Formicinae*^P^, *Kyromyrma* Grimaldi and Agosti, 2000.

#### Discussion

*Kyromyrma* has a generalised morphology (Grimaldi and Agosti 2000) and thus cannot belong to any recent tribe. It has been assumed that *Kyromyrma* is a representative of stem formicines (Ward 2007), i.e., belongs to the clade *Pan-Formicinae*^P^. Given that the crown group *Formicinae*^P^ arose around 80 Ma ago (Brady et al. 2006), the age of *Kyromyrma* (92 Ma) is consistent with this assumption.

### Pan-Dolichoderinae

#### Diagnosis (workers, gynes)

(1) junction of pygidium and hypopygium slit-like (apomorphy); (2) sting vestigial; (3) helcium attached low on anterior face of first gastral segment.

#### Comment

Under the PhyloCode, the taxon should be named “*Pan-Dolichoderinae*^P^, new clade name” and defined as “the total clade composed of the crown clade *Dolichoderinae*^P^ and all extinct species that share a more recent common ancestor with *Dolichoderinae*^P^ than with any extant species that are not members of *Dolichoderinae*^P^”.

#### Composition

*Dolichoderinae*^P^*, Eotapinoma* Dlussky, 1988, *Zherichinius* Dlussky, 1988, *Chronomyrmex* McKellar, Glasier and Engel, 2013.

#### Discussion

*Eotapinoma* (Sakhalin and Canadian amber) looks much like representatives of the tribe Tapinomini; but the species from Sakhalin amber (Dlussky 1988) are poorly preserved, and the one from Canadian amber (Dlussky 1999a) was described briefly and is now lost. Dlussky also noted that *Eotapinoma* and *Zherichinius* (Sakhalin amber) have similarities with both the Dolichoderinae and Formicinae (Dlussky 1988, 1999a). However, there is now little doubt that both genera are stem dolichoderines (Ward et al. 2010). One cannot also reject the possibility that *Zherichinius* is a crown dolichoderine (*Dolichoderinae*^P^), since the latter arose around 60-67 Ma ago (Ward et al. 2010) and thus can in principle be present in Sakhalin amber, which is 43-47 Ma old (Radchenko and Perkovsky 2016).

*Chronomyrmex* (Canadian amber) initially was placed in the tribe Leptomyrmecini (McKellar et al. 2013a). However, the Leptomyrmecini is a morphologically heterogeneous assemblage, recognized primarily by disagreement with the three other tribes (Ward et al. 2010), and thus it is obvious that *Chronomyrmex* simply lacks the characters of the other tribes. Taking into account the time of its emergence, *Chronomyrmex* cannot belong to the crown dolichoderines (and as a result, to any recent dolichoderine tribe); it is a stem taxon to the Dolichoderinae, or, less probably, to all dolichoderomorphs (Dolichoderinae + Aneuretinae).

### Pan-Ectatomminae

#### Diagnosis (workers, gynes)

(1) clypeus broadly inserted between frontal lobes; (2) outer margins of frontal lobes not pinched in posteriorly; (3) helcium projects from about midheight of anterior face of abdominal segment III; no high vertical anterior face to abdominal segment III above helcium.

#### Comment

Under the PhyloCode, the taxon should be named “*Pan-Ectatomminae*^P^, new clade name” and defined as “the total clade composed of the crown clade *Ectatomminae*^P^ and all extinct species that share a more recent common ancestor with *Ectatomminae*^P^ than with any extant species that are not members of *Ectatomminae*^P^”.

#### Composition

*Ectatomminae*^P^, *Canapone* Dlussky, 1999.

#### Discussion

The first and second diagnostic characters are putative, as the morphology of the anterodorsal part of the head in *Canapone* (Canadian amber) is unknown, and the holotype is now lost. *Canapone* initially was placed in the Ponerinae (Dlussky 1999a) and then transferred to the Ectatomminae incertae sedis (Bolton 2003). It is likely that *Canapone* is closest to the Ectatomminae, but cannot be placed in any recent ectatommine tribe, as it is unique in having plesiomorphies that have been lost by extant species (Bolton 2003).

### Aculeata incertae sedis

#### Genera

*Cretopone* Dlussky, 1975, *Dolichomyrma* Dlussky, 1975, *Khetania* Dlussky, 1999, *Petropone* Dlussky, 1975.

#### Discussion

*Cretopone*, *Petropone*, and *Khetania* are poorly preserved. It is hard to disagree with Grimaldi et al.’s (1997) conclusion that the first two genera do not have ant synapomorphies. *Khetania* does not have ant synapomorphies either; its petiole is not well defined, the antennae are not preserved. *Dolichomyrma* is remarkable for its small size (3-5 mm) and the absence of wings, as in worker ants. The petiole of *Dolichomyrma* is nodeless, which is why Dlussky initially believed it was a dolichoderine or specialized sphecomyrmine (Dlussky 1975), but then placed it in the Armaniidae (Dlussky 1983). I am proposing that *Dolichomyrma* be placed into the Aculeata incertae sedis, because it does not have ant synapomorphies, and its petiole is similar to that of some Bethylidae.

### Conclusion: the origin and evolution of ants

Fifty years ago, Wilson et al. (1967) discovered Cretaceous *Sphecomyrma,* a primitive ant with plesiomorphic characters, claimed to be the ancestor either of one of the two branches of the ant lineage (Wilson et al. 1967) or all living ants (Taylor 1978). Since then, other ants, some 10 Ma older than *Sphecomyrma,* have been discovered. It has become evident that Cretaceous stem-group ants were not only very diverse but also very specialized. Furthermore, if primitive stem lineages had coexisted with crown-group ants, such as *Kyromyrma*, *Canapone*, *Eotapinoma*, *Brownimecia,* then the former cannot be the direct ancestors of the latter. There is now general agreement that stem groups like the Sphecomyrminae are the result of the primary diversification in the ant tree, and so the true ancestor of both stem- and crown-group ants has to have existed before them.

Currently, however, there is no consensus of opinion on what that ancestor was like. Wheeler (1926) proposed that the genus *Myzinum* (Tiphiidae) is closest to the ants; Wilson et al. (1967) concurred about Tiphiidae but chose the genus *Methocha* Latreille. Dlussky and Fedoseeva (1988) argued that groups with wingless females cannot be ant ancestors because this leaves unexplained the secondary emergence of wings in ants. After the rise of cladistics and introduction of the methods of molecular systematics, the issue became no less clear. The first morphological cladistic study by Brothers (1999) showed the sister group of ants to be Vespidae + Scoliidae. A DNA study of the Hymenoptera (Heraty et al. 2011) suggested that the sister groups of ants is either Mutillidae + Sapygidae + Tiphiidae + Bradynobaenidae + Pompilidae + Scoliidae or Sphecidae + Scoliidae. Another study (Peters et al. 2011) revealed it to be either Vespidae + Mutillidae + Bradynobaenidae + Bethylidae + Pompilidae or Tiphidae. A study combining molecular and morphological data of the Vespoidea (Pilgrim et al. 2008) showed it is either Sapygidae + Bradynobaenidae or Vespidae + Rhopalostomatidae. Finally, a phylogenomic study (Johnson et al. 2013) based on the genomes and transcriptomes of 11 species of the Aculeata unexpectedly concluded that ants are closer to the Apoidea, not to Vespoidea.

Most of the presently known stem-group ants are thought to have had an arboreal lifestyle - they have long legs and are preserved in amber, ancient tree resin. On the other hand, most primitive extant ants (Martialinae, Leptanillinae, poneroids) are small cryptic subterranean species. These groups could indeed have evolved from above-ground ancestors, but since the general trend of ant morphological evolution suggests otherwise, there is some reason to think that their ancestors were cryptic as well. The increased mobility of the gaster, with the resulting separation of the petiole, likely suggests an adaptation to an underground lifestyle. This type of adaptation is also recognizable in the emergence of the metapleural gland, which has a role in defense against parasites in underground colonies (Yek and Mueller 2011). Therefore, if the first ants were underground (Lucky et al. 2013), then the Martialinae and other primitive forms may be viewed as relicts that have changed little during evolution as a result of living in ecologically stable habitats (Rabeling et al. 2008).

Finding paleontological records of the subterranean ant ancestors is a significant challenge. The reasons are: (1) that these ants are hardly preserved in amber due to their cryptic lifestyle (for example, ants are unknown in Early Cretaceous ambers (LaPolla et al. 2013) such as Spanish and Lebanese), (2) that their number was too small to be occasionally trapped in amber (ants comprise only 0.001 - 0.05% of all insects preserved in Late Cretaceous ambers (Grimaldi and Agosti 2000), so the number of ants in earlier ambers is even lower), and (3) that they were too small to be preserved as imprints in rock (however, ichnofossils, such as those from the Upper Jurassic Morrison Formation dated at 156-146 Ma (Hasiotis and Demko 1996), may be traces of these ants).

When considering factors underlying ants ‘ extraordinary evolutionary success, phenotypic plasticity and ecological niche construction have to be named first. The former is the capacity of a single genotype to exhibit variable phenotypes - behavioral, biochemical, physiological, developmental - in different environments (West-Eberhard 2003; Pigliucci et al. 2006). It is now viewed as a widespread phenomenon that can facilitate evolutionary change and speciation (Price et al. 2003). One example of ants’ phenotypic plasticity in action is caste polyphenism responsible for their diverse ecological adaptation (Simpson et al. 2011). The second factor, niche construction, signifies the alteration of the environment that then affects selection pressures. Odling-Smee et al. (2003) claimed that niche construction “should be regarded, after natural selection, as a second major participant in evolution”. The importance of ant niche construction is difficult to overestimate, as ants are among the most active ecosystem engineers (Folgarait 1998).

Given the aforementioned ideas, the main steps in ant evolution can be now outlined. The first ants, which may have originated as early as the Upper Jurassic, were solitary underground species. During the Late Cretaceous, about 100 Ma ago, they underwent diversification, evolved a nuptial flight and arboreal lifestyle, either become eusocial or were at the stage of facultative sociality. The common ancestor of crown-group ants lived about 123 Ma (from 141 to 116 Ma) ago (Brady et al. 2006; Schmidt 2013); thus stem- and crown-group ants had existed alongside one another throughout the Cretaceous period, undergoing spectacular speciation in the first angiosperm forests (Moreau et al. 2006).

There was the transformation of terrestrial ecosystems after the major biotic extinction event at the end of the Mesozoic, during which about 50% of genera and 75% of plant and animal species became extinct (Jablonski and Chaloner 1994). Arboreal species were probably most vulnerable at that time and thereby doomed. Only some ants that now compose the crown clade have survived and successfully crossed the K/Pg boundary. They then occupied vacant niches of above-ground and arboreal predators and also began to actively make new ecological niches, thus preparing their own huge evolutionary success.

From this view, the evolutionary destiny of ants is similar to the one of mammals which occupied new niches after the extinction of large reptiles. During Cenozoic time, both groups have undergone remarkable adaptive radiation; in the invertebrate micro-world and vertebrate macro-world respectively, they became successful terrestrial predators, largely thanks to phenotypic plasticity, brood care and complex social behavior.

## Acknowledgments

I thank A. Romain and J. Glasier for improving a draft of the manuscript, as well as R. Kaur, O. Nordmann and A. Radchenko for their valuable comments.

## Literature Cited

Agosti, D., J. Moog, and U. Maschwitz. 1999. Revision of the Oriental plant-ant genus *Cladomyrma*. American Museum Novitates 3283: 1–24.

Alpert, G. D. 2007. A review of the ant genus *Metapone* Forel from Madagascar. Memoirs of the American Entomological Institute 80: 8–18.

Archibald, S. B., K. R. Johnson, R. W. Mathewes, and D. R. Greenwood. 2011. Intercontinental dispersal of giant thermophilic ants across the Arctic during early Eocene hyperthermals. Proceedings of the Royal Society (Series B, Biological Sciences) 278: 3679–3686.

Ax, P. 1985. Stem species and the stem lineage concept. Cladistics 1: 279–287.

Baroni Urbani, C., B. Bolton, and P. S. Ward. 1992. The internal phylogeny of ants (Hymenoptera: Formicidae). Systematic Entomology 17: 301–329.

Baroni Urbani, C., and M. L. de Andrade. 2003. The ant genus *Proceratium* in the extant and fossil record (Hymenoptera: Formicidae). Regionale di Scienze Naturali Monografie (Turin) 36: 1–492.

Barden, P., and D. A. Grimaldi. 2012. Rediscovery of the bizarre Cretaceous ant *Haidomyrmex* Dlussky (Hymenoptera: Formicidae), with two new species. American Museum Novitates 3755: 1–16.

Barden, P., and D. Grimaldi. 2013. A new genus of highly specialized ants in Cretaceous Burmese amber (Hymenoptera: Formicidae). Zootaxa 3681: 405–412.

Barden, P., and D. Grimaldi. 2014. A Diverse ant fauna from the Mid-Cretaceous of Myanmar (Hymenoptera: Formicidae). PloS One 9: e93627.

Barden, P., and D. Grimaldi. 2016. Adaptive radiation in socially advanced stem-group ants from the Cretaceous. Current Biology 26: 515–521.

Baum, D. A., and S. D. Smith. 2012. Tree thinking: an introduction to phylogenetic biology. Roberts and Company; Greenwood Village, CO. 476 p.

Bolton, B. 1973. A remarkable new arboreal ant genus (Hym. Formicidae) from West Africa. Entomologist’s Monthly Magazine 108: 234–237.

Bolton, B. 1994. Identification guide to the ant genera of the world. Harvard University Press; Cambridge. 222 p.

Bolton, B. 2003. Synopsis and classification of Formicidae. Memoirs of the American Entomological Institute 71: 1–370.

Bolton, B. 2007. Taxonomy of the dolichoderine ant genus *Technomyrmex* Mayr (Hymenoptera: Formicidae) based on the worker caste. Contributions of the American Entomological Institute 35: 1–150.

Bolton, B., and B. L. Fisher. 2008. Afrotropical ants of the ponerine genera *Centromyrmex* Mayr, *Promyopias* Santschi gen. rev. and *Feroponera* gen. n., with a revised key to genera of African Ponerinae (Hymenoptera: Formicidae). Zootaxa 1929: 1–37.

Bolton, B., and B. L. Fisher. 2011. Taxonomy of Afrotropical and West Palaearctic ants of the ponerine genus *Hypoponera* Santschi (Hymenoptera: Formicidae). Zootaxa 2843: 1–118.

Boudinot, B. E. 2015. Contributions to the knowledge of Formicidae (Hymenoptera, Aculeata): a new diagnosis of the family, the first global male-based key to subfamilies, and a treatment of early branching lineages. European Journal of Taxonomy 120: 1–62.

Brady, S. G., B. L. Fisher, T. R. Schultz, and P. S. Ward. 2014. The rise of army ants and their relatives: diversification of specialized predatory doryline ants. BMC Evolutionary Biology 14: 93.

Brady, S. G., T. R. Schultz, B. L. Fisher, and P. S. Ward. 2006. Evaluating alternative hypotheses for the early evolution and diversification of ants. Proceedings of the National Academy of Sciences 103: 18172–18177.

Brady, S. G., and P. S. Ward, P.S. 2005. Morphological phylogeny of army ants and other dorylomorphs (Hymenoptera: Formicidae). Systematic Entomology 30: 593–618.

Brothers, D. J. 1999. Phylogeny and evolution of wasps, ants and bees (Hymenoptera, Chrysidoidea, Vespoidea and Apoidea). Zoologica Scripta 28: 233–250.

Cantino, P. D., and K. de Queiroz. 2010. International code of phylogenetic nomenclature. (Available at ∼ http://www.ohiou.edu/phylocode/. Last accessed March 2017.)

Dlussky, G. M. 1975. Superfamily Formicoidea Latreille, 1802. Family Formicidae Latreille, 1802. p. 114–122. *In:* A.P. Rasnitsyn (ed.). Hymenoptera Apocrita of Mesozoic. Transactions of the Paleontological Institute, Academy of Sciences of the Union of Soviet Socialist Republics 147: 114–122. [In Russian].

Dlussky, G. M. 1983. A new family of Upper Cretaceous Hymenoptera: an “intermediate link” between the ants and the scolioids. Paleontologicheskii Zhurnal 3: 65–78. [In Russian].

Dlussky, G. M. 1987. New Formicoidea (Hymenoptera) of the Upper Cretaceous. Paleontologicheskii Zhurnal 1: 131–135. [In Russian.]

Dlussky, G. M. 1988. Ants of Sakhalin amber (Paleocene?). Paleontologicheskii Zhurnal 1: 50–61. [In Russian.]

Dlussky, G. M. 1996. Ants (Hymenoptera: Formicidae) from Burmese amber. Paleontological Journal 30: 449–454.

Dlussky, G. M. 1999a. New ants (Hymenoptera, Formicidae) from Canadian amber. Paleontologicheskii Zhurnal 4: 73–76. [In Russian.]

Dlussky, G. M. 1999b. The first find of the Formicoidea (Hymenoptera) in the lower Cretaceous of the northern hemisphere. Paleontologicheskii Zhurnal 3: 62–66. [In Russian.]

Dlussky, G. M., D. J. Brothers, and A. P. Rasnitsyn. 2004. The first Late Cretaceous ants (Hymenoptera: Formicidae) from southern Africa, with comments on the origin of the Myrmicinae. Insect Systematics and Evolution 35: 1–13.

Dlussky, G. M., and E. B. Fedoseeva. 1988. Origin and early stages of evolution in ants. p. 70–144. *In:* A. G. Ponomarenko (ed.). Cretaceous biocenotic crisis and insect evolution. Nauka; Moscow. 230 p. [In Russian.]

Dubois, A. 2007. Phylogeny, taxonomy and nomenclature: the problem of taxonomic categories and of nomenclatural ranks. Zootaxa 1519: 27–68.

Engel, M.S., and D. A. Grimaldi. 2005. Primitive new ants in Cretaceous amber from Myanmar, New Jersey, and Canada (Hymenoptera: Formicidae). American Museum Novitates 3485: 1–23.

Ereshefsky, M. 2001. The poverty of the Linnaean hierarchy: a philosophical study of biological taxonomy. Cambridge University Press; Cambridge, U.K. 316 p.

Faul, F., E. Erdfelder, A.-G. Lang, and A. Buchner. 2007. G*Power 3: A flexible statistical power analysis program for the social, behavioral, and biomedical sciences. Behavior Research Methods 39: 175–191.

Fisher, B. L. 2002. AntWeb. (Available at ∼ http://www.antweb.org. Last accessed March 2017.)

Folgarait, P. J. 1998. Ant biodiversity and its relationship to ecosystem functioning: a review. Biodiversity and Conservation7. 9: 1221–1244.

Greggers, U., G. Koch, V. Schmidt, A. Dürr, A. Floriou-Servou, D. Piepenbrock, M. C. Göpfert, and R. Menzel R. 2013. Reception and learning of electric fields in bees. Proceedings of the Royal Society B 280: doi:10.1098/rspb.2013.0528.

Grimaldi, D., and D. Agosti, D. 2000. A formicine in New Jersey Cretaceous amber (Hymenoptera: Formicidae) and early evolution of the ants. Proceedings of the National Academy of Sciences of the United States of America 97: 13678–13683.

Grimaldi, D., D. Agosti, and J. M. Carpenter. 1997. New and rediscovered primitive ants (Hymenoptera: Formicidae) in Cretaceous amber from New Jersey, and their phylogenetic relationships. American Museum Novitates 3208: 1–43.

Hasiotis S.T., and T. M. Demko. 1996. Terrestrial and freshwater trace fossils, upper Jurassic Morrison formation, Colorado Plateau. p. 355–370. *In:* M. Morales (Ed.) The Continental Jurassic. Museum of Northern Arizona Bulletin, 60: 355–370.

Hennig, W. 1966. Phylogenetic systematics. University of Illinois Press; Urbana, IL. 263 p.

Heraty, J., F. Ronquist, J. M. Carpenter, D. Hawks, S. Schulmeister, A. P. Dowling, et al. 2011. Evolution of the hymenopteran megaradiation. Molecular Phylogenetics and Evolution 60: 73–88.

Hölldobler, B., and E. O. Wilson. 1990. The ants. Belknap Press; Cambridge, MA. 732 p.

Jablonski, D., and W. G. Chaloner. 1994. Extinctions in the fossil record (and discussion). Philosophical Transactions of the Royal Society of London, Series B 344: 11–17.

Johnson, B. R., M. L. Borowiec, J. C. Chiu, E. K. Lee, J. Atallah, and P. S. Ward. 2013. Phylogenomics resolves evolutionary relationships among ants, bees, and wasps. Current Biology 23: 2058–2062.

Joyce, W. G., Parham J. F., and Gauthier J. A. 2004. Developing a protocol for the conversion of rank-based taxon names to phylogenetically defined clade names, as exemplified by turtles. Journal of Paleontology 78: 989–1013.

Kamikouchi, A., H. K. Inagaki, T. Effertz, O. Hendrich, A. Fiala, M. C. Göpfert, and K. Ito. 2009. The neural basis of *Drosophila* gravity-sensing and hearing. Nature 458: 165–171.

LaPolla, J. S. 2009. Taxonomic revision of the Southeast Asian ant genus *Euprenolepis*. Zootaxa 2046: 1–25.

LaPolla, J. S., G. M. Dlussky, and V. Perrichot. 2013. Ants and the Fossil Record. Annual review of Entomology 58: 609–630.

Linnaeus, C. 1761. Fauna suecica sistens animalia Sueciae regni: Mammalia, Aves, Amphibia, Pisces, Insecta, Vermes. L. Salvii; Stockholm. 578 p.

Lucky, A., M. D. Trautwein, B. S. Guenard, M. D. Weiser, and R. R. Dunn. 2013. Tracing the rise of ants-out of the ground. PloS One 8: e84012.

Lucky, A., and P. S. Ward. 2010. Taxonomic revision of the ant genus *Leptomyrmex* Mayr (Hymenoptera: Formicidae). Zootaxa 2688: 1–67.

MacKay, W. P. 1993. A review of the new world ant of the genus *Dolichoderus*. Sociobiology 22: 1–148.

McDonald, J. H. 2014. Handbook of Biological Statistics (3rd ed.). Sparky House Publishing; Baltimore, MD. 299 p.

McKellar, R. C., J. R. N. Glasier, and M. S. Engel. 2013a. New ants (Hymenoptera: Formicidae: Dolichoderinae) from Canadian Late Cretaceous amber. Bulletin of Geosciences 88: 583–594.

McKellar, R. C., Glasier, J. R. N. Glasier, and M. S. Engel. 2013b. A new trap-jawed ant (Hymenoptera: Formicidae: Haidomyrmecini) from Canadian Late Cretaceous amber. The Canadian Entomologist 145: 454–465.

McKellar, R. C. and A. P. Wolfe. 2010. Canadian amber. p. 96–113. *In:* D. Penney (ed.). Biodiversity of fossils in amber from the major world deposits. Siri Scientific Press; Manchester. 304 p.

Moreau, C. S., C. D. Bell, R. Vila, S. B. Archibald, and N. E. Pierce. 2006. Phylogeny of the ants: diversification in the age of Angiosperms. Science 312: 101–104.

Nel, A., G. Perrault, V. Perrichot, and D. Néraudeau. 2004. The oldest ant in the Lower Cretaceous amber of Charente-Maritime (SW France) (Insecta: Hymenoptera: Formicidae). Geologica Acta 2: 23–29.

Odling-Smee, F. J., K. N. Laland, and M. W. Feldman. 2003. Niche construction: the neglected process in evolution. Princeton University Press; Princeton, NJ. 472 p.

Perrichot, V. 2014. A new species of the Cretaceous ant *Zigrasimecia* based on the worker caste reveals placement of the genus in the Sphecomyrminae (Hymenoptera: Formicidae). Myrmecological News 19: 165–169.

Perrichot, V. 2015. A new species of *Baikuris* (Hymenoptera: Formicidae: Sphecomyrminae) in mid-Cretaceous amber from France. Cretaceous Research 52: 585–590.

Perrichot, V., A. Nel, D. Neraudeau, S. Lacau, and T. Guyot. 2008. New fossil ants in French Cretaceous amber (Hymenoptera: Formicidae). Naturwissenschaften 95: 91–97.

Perrichot, V., B. Wang, and M. S. Engel. 2016. Extreme morphogenesis and ecological specialization among Cretaceous basal ants. Current Biology 26: 1468–1472.

Peters, R. S., B. Meyer, L. Krogmann, J. Borner, K. Meusemann, K. Schütte, et al. 2011. The taming of an impossible child: a standardized all-in approach to the phylogeny of Hymenoptera using public database sequences. BMC Biology 9: 55.

Pigliucci, M., C. J. Murren, and C. D. Schlichting. 2006. Phenotypic plasticity and evolution by genetic assimilation. Journal of Experimental Biology 209: 2362–2367.

Pilgrim, E. M., C. D. Von Dohlen, and J. P. Pitts. 2008. Molecular phylogenetics of Vespoidea indicate paraphyly of the superfamily and novel relationships of its component families and subfamilies. Zoologica Scripta 37: 539–560.

Pleijel, F., and G. W. Rouse. 2003. Cecin’est pas une pipe: names, clades and phylogenetic nomenclature. Journal of Zoological Systematics and Evolutionary Research 41: 162–174.

Price, T. D., A. Qvarnström, and D. E. Irwin. 2003. The role of phenotypic plasticity in driving genetic evolution. Proceedings of the Royal Society of London (Series B: Biological Sciences) 270: 1433–1440.

Rabeling, C., J. M. Brown, and M. Verhaagh. 2008. Newly discovered sister lineage sheds light on early ant evolution. Proceedings of the National Academy of Sciences 105: 14913–14917.

Radchenko, A. G. 1991. Ants of the genus *Strongylognathus* (Hymenoptera, Formicidae) of the fauna of the USSR. Zoologicheskii Zhurnal 70: 84–90. [in Russian]

Radchenko, A. G. 1994. Identification table for ants of the genus *Myrmica* (Hymenoptera, Formicidae) from central and eastern Palearctic. Zoologicheskii Zhurnal 73: 130–145. [In Russian]

Radchenko, A. G., and G. W. Elmes. 1998. Taxonomic revision of the ritae species-group of the genus *Myrmica* (Hymenoptera, Formicidae). Vestnik Zoologii 32: 3–27.

Radchenko, A., and G. W. Elmes. 2003. A taxonomic revision of the socially parasitic *Myrmica* ants (Hymenoptera: Formicidae) of the Palaearctic region. Annales Zoologici 53: 217–243.

Radchenko, A. G., and E. E. Perkovsky. 2016. The ant *Aphaenogaster dlusskyana* sp. nov. (Hymenoptera, Formicidae) from the Sakhalin amber – the earliest described species of an extant genus of Myrmicinae. Paleontological Journal 50: 936–946.

Radchenko, A., G. W. Elmes, and M. Woyciechowski. 2002. An appraisal of *Myrmica bergi* Ruzsky, 1902 and related species (Hymenoptera: Formicidae). Annales Zoologici (Warsawa) 52: 409–421.

Schmidt, C. 2013. Molecular phylogenetics of ponerine ants (Hymenoptera: Formicidae: Ponerinae). Zootaxa 3647: 201–250.

Schmidt, F. A., Feitosa, R. M., de Moraes Rezende, F., and Silva de Jesus, R. (2013) News on the enigmatic ant genus *Anillidris* (Hymenoptera: Formicidae: Dolichoderinae: Leptomyrmecini). Myrmecological News 19: 25–30.

Seifert, B. 1988. A taxonomic revision of the *Myrmica* species of Europe, Asia Minor, and Caucasus (Hymenoptera, Formicidae). Abhandlungen und Berichte des Naturkundemuseums Görlitz 62: 1–75.

Seifert, B. 1992. A taxonomic revision of the Palaearctic members of the ant subgenus *Lasius* s. str.(Hymenoptera, Formicidae). Abhandlungen und Berichte des naturkundemuseums Görlitz 66: 1–67.

Seifert, B. 2000. A taxonomic revision of the ant subgenus *Coptoformica* Mueller, 1923 (Hymenoptera, Formicidae). Zoosystema-Paris 22: 517–568.

Seifert, B. 2003. The ant genus *Cardiocondyla* (Insecta: Hymenoptera: Formicidae) - a taxonomic revision of the *C. elegans, C. bulgarica, C. batesii, C. nuda, C. shuckardi, C. stambuloffii, C. wroughtonii, C. emeryi* and *C. minutior* species groups. Annalen des Naturhistorischen Museums in Wien (B, Botanik, Zoologie) 104: 203–338.

Shattuck, S. O. 1992. Generic revision of the ant subfamily Dolichoderinae (Hymenoptera: Formicidae). Sociobiology 21: 1–181.

Shattuck, S. O. 2008. Australian ants of the genus *Aphaenogaster* (Hymenoptera: Formicidae). Zootaxa 1677: 25–45.

Shattuck, S. O., and N. J. Barnett. 2006. Australian species of the ant genus *Diacamma* (Hymenoptera: Formicidae). Myrmecologische Nachrichten 8: 13–19.

Simpson, S. J., G. A. Sword, and N. Lo. 2011. Polyphenism in insects. Current Biology 21: R738–R749.

Stephens, J. F. 1829. A systematic catalogue of British insects: being an attempt to arrange all the hitherto discovered indigenous insects in accordance with their natural affinities. Baldwin and Cradock; London. 388 p.

Taylor, R. W. 1978. *Nothomyrmecia macrops*: a living-fossil ant rediscovered. Science 201: 979–985.

Taylor, R. W. 1980. Notes on the Russian endemic ant genus *Aulacopone* Arnoldi (Hymenoptera: Formicidae). Psyche (Cambridge) 86: 53–361.

Ward, P. S. 1999. Systematics, biogeography and host plant associations of the *Pseudomyrmex viduus* group (Hymenoptera: Formicidae), Triplaris-and Tachigali-inhabiting ants. Zoological Journal of the Linnean Society 126: 451–540.

Ward, P. S. 2007. Phylogeny, classification, and species-level taxonomy of ants (Hymenoptera: Formicidae). Zootaxa 1668: 549–563.

Ward, P. S. 2011. Integrating molecular phylogenetic results into ant taxonomy (Hymenoptera: Formicidae). Myrmecological News 15: 21–29.

Ward, P. S., S. G. Brady, B. L. Fisher, and T. R. Schultz. 2010. Phylogeny and biogeography of dolichoderine ants: effects of data partitioning and relict taxa on historical inference. Systematic Biology 59: 342–362.

West-Eberhard, M. J. 2003. Developmental plasticity and evolution. Oxford University Press; New York, NY. 794 p.

Wheeler, W. M. 1926. Les sociétés d’insectes: leur origine, leur évolution. Gaston Doin and Co.; Paris, France. 468 p.

Wild, A. L. 2004. Taxonomy and distribution of the Argentine ant, *Linepithema humile* (Hymenoptera: Formicidae). Annals of the Entomological Society of America 97: 1204–1215.

Wilson, E. O. 1985. Ants from the Cretaceous and Eocene amber of North America. Psyche 92: 205–216.

Wilson, E. O. 1987. The earliest known ants: an analysis of the Cretaceous species and an inference concerning their social organization. Paleobiology 13: 44–53.

Wilson, E. O., F. M. Carpenter, and W. L. Brown. 1967. The first Mesozoic ants, with the description of a new subfamily. Psyche 74: 1–19.

Wilson, E.O., and B. Hölldobler. 2005. The rise of the ants: a phylogenetic and ecological explanation. Proceedings of the National Academy of Sciences 102: 7411–7414.

Yanega, D. 1997. Demography and sociality in halictine bees (Hymenoptera: Halictidae). p. 293–315. *In:* J. Choe, and B. Crespi (eds.). The evolution of social behavior in insects and arachnids. Cambridge University Press; Cambridge, U.K. 552 p.

Yek, S. H., and U. G. Mueller. 2011. The metapleural gland of ants. Biological Reviews 86: 774–791.

